# Electrostatic control of conformational dynamics drives proton translocation in respiratory Complex I

**DOI:** 10.64898/2026.11.03.764137

**Authors:** Jobst Liebau, Patricia Saura, Sofia Badolato, Ana P. Gamiz-Hernandez, Ville R. I. Kaila

**Author notes:** **Corresponding author:** Ville R. I. Kaila. equal contributions (Jobst Liebau, Patricia Saura).

## Abstract

Complex I is a large membrane protein that transduces redox energy into a proton motive force across a biological membrane, powering cellular respiration. Yet, despite significant experimental and computational efforts, its long-range proton translocation mechanism remains poorly understood. Here, we combine nuclear magnetic resonance spectroscopy and site-directed mutagenesis experiments with multiscale classical and hybrid quantum/classical (QM/MM) simulations to probe how conformational dynamics is linked to proton transfer reactions in the terminal antiporter module (Nqo12/NuoL/ND5) of Complex I. We show that protonation of a conserved proton-loading site (Lys329/His241^Nqo12^) in the terminal antiporter-like module gives rise to conformational changes that result in opening of the proton pathway. We further show that this protonation reaction is controlled by buried ion-pairs that modulate the barrier for proton translocation by intrinsic electric field effects. On a general level, our combined findings reveal how bioenergetic machines combine conformational dynamics with hydration transitions to establish efficient redox-driven proton pumps.

## INTRODUCTION

Complex I (NADH:quinone oxidoreductase) is a 0.5-1 MDa membrane-bound protein complex that powers oxidative phosphorylation in mitochondria and many bacteria (*1*). Complex I transduces redox energy by transferring electrons from nicotinamide adenine dinucleotide (NADH) to quinone (Q) in its hydrophilic domain, which couples to proton pumping across its 200 Å wide membrane domain, generating a proton motive force (PMF) that drives the synthesis of adenosine triphosphate (ATP). The redox reactions involve a 100 Å long electron transfer process, catalyzed by 8 iron-sulfur (FeS) centers, which provide a tunneling wire for the Q reduction process (*2*, *3*), while the proton pumping activity is achieved by the antiporter-like Nqo12, Nqo13, and Nqo14 modules (in *T. thermophilus* Complex I, NuoL/NuoM/NuoN in *E. coli* Complex I; ND5/ND4/ND2 in mitochondrial Complex I), together with Nqo7/10/11/8 (NuoA/NuoJ/NuoK/NuoH; ND3/ND6/ND4L/ND1) (*4–6*), located at the interface of the hydrophilic and membrane domains (Fig. 1A). This long-range proton-coupled electron transfer (PCET) process is highly efficient and fully reversible, but its mechanistic principles remain poorly understood, with several competing models put forward in recent years (*7–13*).

**Figure 1.**
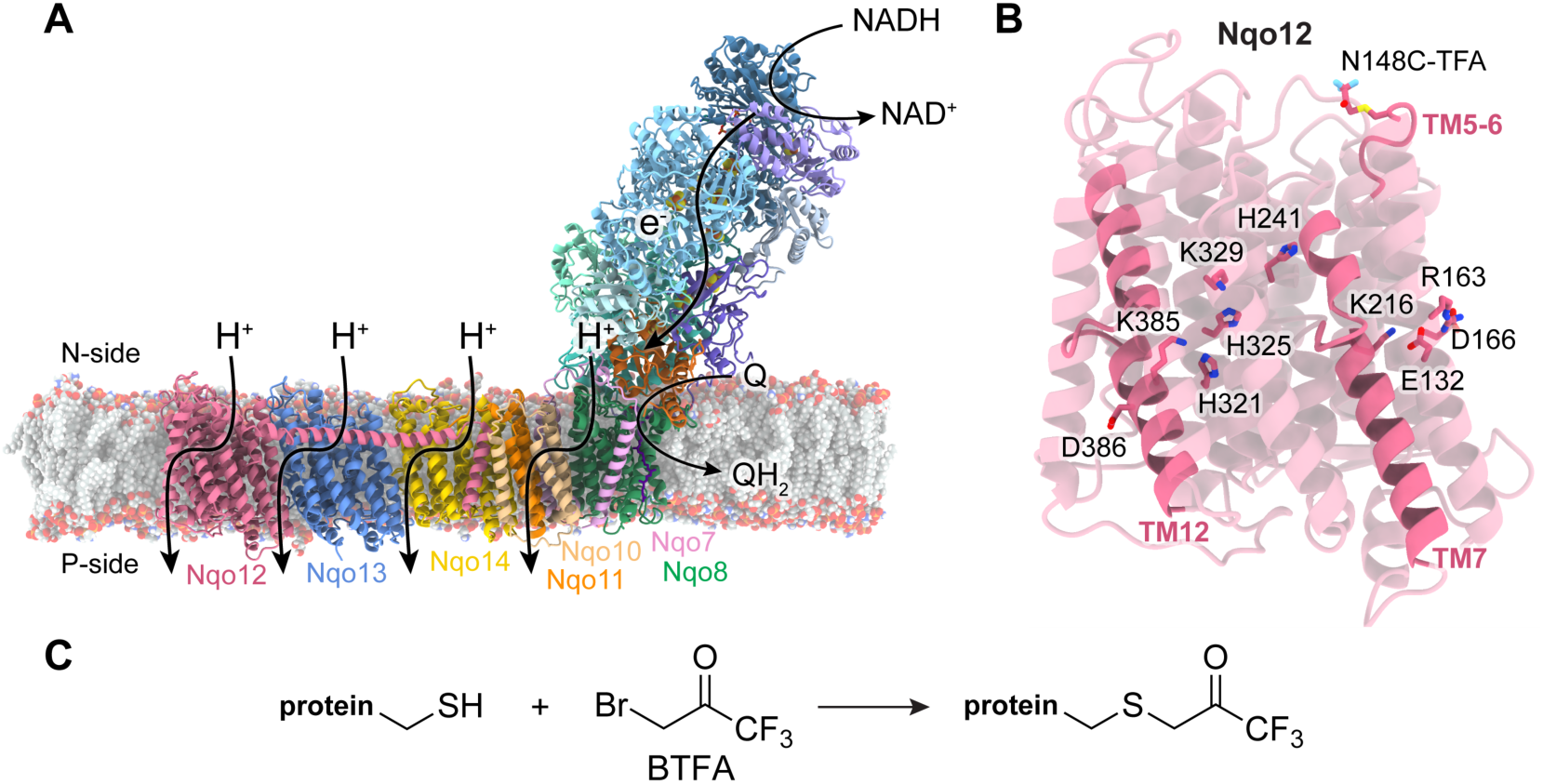
Structure and function of Complex I and the terminal Nqo12 antiporter-like module. (**A**) NADH-driven quinone reduction drives proton pumping across the membrane domain. (**B**) Structure of Nqo12, functional residues along the proton pathway, and the position of the NMR label. (**C**) ^19^F-labeling of a cysteine residue with 3-bromo-1,1,1-trifluoroacetone (BTFA), resulting in a trifluoro-acetone (TFA) cysteine.

The tightly coupled PCET reactions in Complex I have been difficult to characterize, as mutations of key residues result in the loss of both the oxidoreductive and proton pumping activities. We previously showed (*14*) that engineered variants of the individual antiporter-like subunits (Nqo12, Nqo13) of Complex I contain all functional elements required to catalyze proton transport across proteoliposome membranes (*cf.* also Refs. (*15*, *16*)). More specifically, the conformational state of a conserved ion-pair, together with non-polar (Leu/Phe) gates in both the terminal (Nqo12) and penultimate (Nqo13) antiporter-like modules, control the rate of proton transfer by modulating reaction barriers and the hydration state of the subunits (*14*, *15*). These findings are consistent with earlier observations by Dröse *et al.* (*17*), showing that a Complex I-knockout, with the terminal and penultimate (Nqo12/13) antiporter-like subunits removed, catalyzes proton pumping with a reduced stoichiometry, thus reinforcing that all antiporter subunits are involved in the pumping process. Taken together, these findings challenge recent mechanistic models (*8*, *9*), which suggest that only the terminal subunit (Nqo12) is responsible for pumping all protons in Complex I.

Molecular dynamics (MD) simulations suggest that the hydration and proton transfer reactions in the antiporter-like subunits induce subtle conformational changes in broken helix elements TM7a/b and TM12a/b, surrounding the proton input and output channels (*16*, *18–20*), together with sidechain flips of key residues along the pathway (*15*, *18*, *20–23*) that support the process. Despite challenges to experimentally observe conformational and hydration changes in Complex I, MD simulations thus provide important guiding principles to probe protein dynamics linked to proton transport.

Here, we employ nuclear magnetic resonance (NMR) spectroscopy of site-specific fluorine (^19^F) labeled antiporter-like subunit Nqo12, in combination with multiscale molecular simulations and proton conduction assays, to identify how the protonation dynamics are coupled to protein conformational changes. While NMR is traditionally challenging to apply to large membrane protein complexes, ^19^F-NMR (*24–27*) is emerging as a powerful tool to probe the functional dynamics of membrane proteins.

## RESULTS

### The terminal antiporter-like subunit samples distinct conformational states

To introduce a ^19^F-NMR probe into the terminal Nqo12 antiporter module, we first analyzed the N-side input channel and dynamics of the broken helix TM7a/b based on atomistic MD simulations in combination with sequence analysis to identify possible non-conserved probe sites (Fig. S1A, C). Consistent with previous findings, we observe an S-shaped proton channel (*cf*. also Ref. (*8*, *15*, *18*, *28–30*)) forming around TM7a, which undergoes a subtle conformational shift upon channel hydration (Fig. 2C). We identified a non-conserved surface-exposed Asn148 residue in the TM5-6 loop, and hypothesized that its conformation could be affected by conformational changes in the channel (Fig. 1B, Fig. 2, Fig. S1A, C). Labeling of the resulting Nqo12-N148C variant with 3-bromo-1,1,1-trifluoroacetone (BTFA) (Fig. 1C) forms a trifluoro-acetone (TFA) cysteine, without spurious labeling of other sites, as demonstrated by our NMR experiments (Fig. 2A, Fig. S1, see *Methods*). Moreover, while labeling of other surface-exposed cysteines introduced into Nqo12 is also possible (Fig. S1B), we focused here on the TM5-6 loop labeled constructs to probe specific conformational changes linked to the proton uptake along the N-side channel (see below).

**Figure 2.**
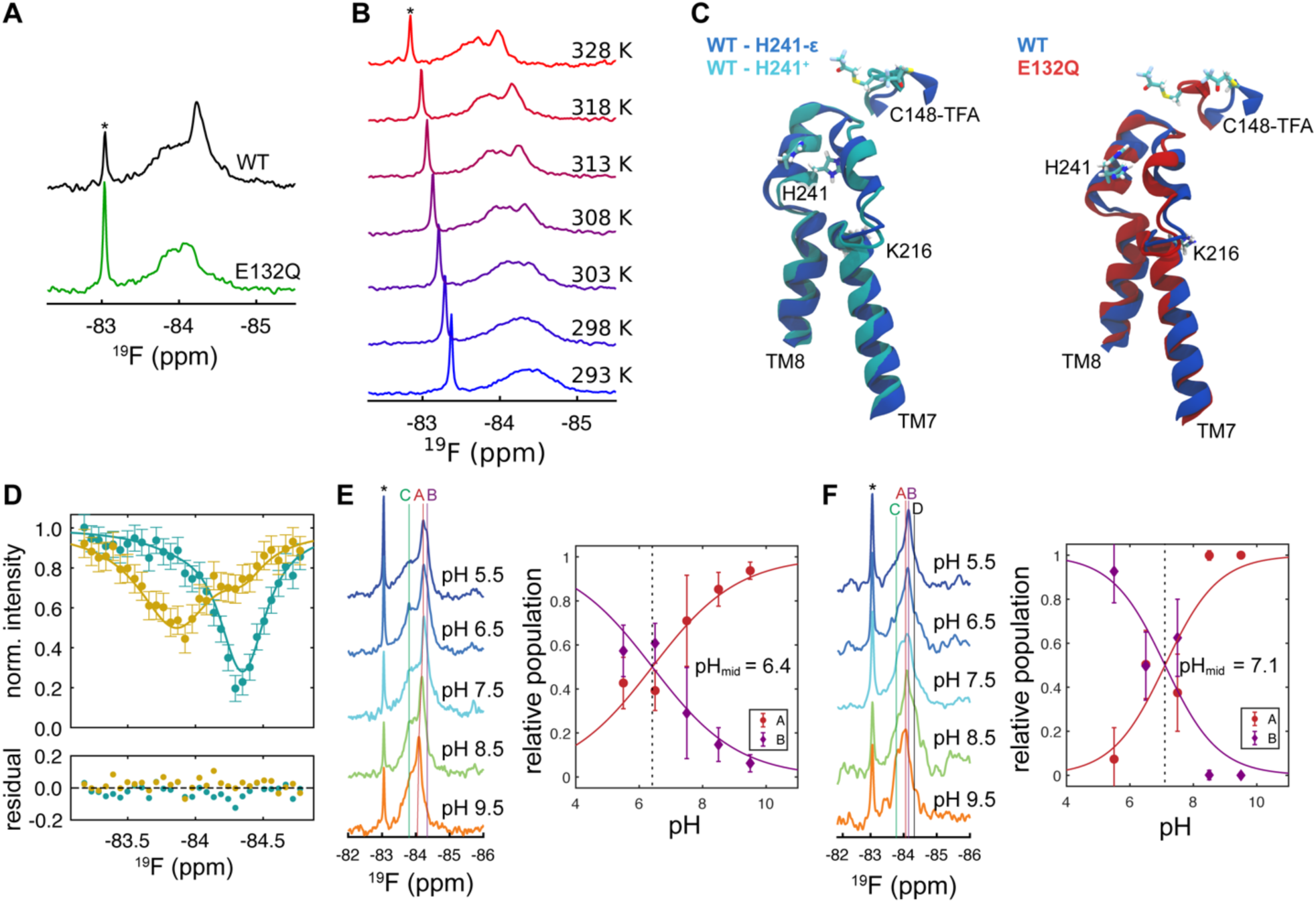
Conformational dynamics of antiporter-like module Nqo12. **(A)** ^19^F-spectra of WT-Nqo12 (black) and E132Q-Nqo12 (green). (**B**) Temperature dependence of WT-Nqo12 ^19^F-spectra. The asterisk indicates the resonance of the free TFA. (**C)** Dynamics of TM7a and TM8 during MD simulations affect the position of the TM5-6 loop, containing the ^19^F-probe. (**D**) ^19^F chemical exchange saturation transfer (CEST) experiment of WT-Nqo12 for resonances **C** (yellow) and **A/B** (teal). The solid lines show a global fit to the data employing a two-state model (see *Methods*), while the lower panel shows the residual of the fit. pH-dependence of the ^19^F-spectra for WT-Nqo12 (**E,** left) and E132Q-Nqo12 (**F,** left). Distinct populations are indicated by vertical lines. Ratio of the pH-dependent resonances **A** and **B** based on line shape fits for (**E,** right) WT, and (**F,** right) E132Q-Nqo12.

At temperatures below 303 K, the Nqo12^N148C-TFA^ variant (from here on, “WT-Nqo12”) shows a broad NMR resonance, assigned here as conformation **C** (Fig. 2B), while with increasing temperature, we resolve two additional resonances upfield from the latter, assigned as conformations **A** and **B**, in which the label samples unique chemical environments. **A** and **B** are narrower than **C** (Fig. 2B), and in contrast to the latter, their line widths are strongly affected by temperature changes, likely arising from increased nanosecond sidechain or loop dynamics (Fig. 2B) (*31*). Our chemical exchange saturation transfer (CEST) experiment shows that the **A** / **B** populations exchange on a seconds timescale (*k*_ex_=1.9±0.8 s^-1^) with **C** (Fig. 2D, Table S2), suggesting that the observed conformations arise from conformational transitions between the substates, thus probing distinct conformations along the proton pathway. Importantly, the relative populations of **A** and **B** show a strong pH-dependence, with the **A** / **B**-ratio titrating with an apparent p*K*_a_ of 6.4 ± 0.2 (Fig. 2E) based on line shape analysis (see *Methods*). Taken together, our data show that the identified states are strongly linked to protonation changes within the antiporter module.

### Protonation dynamics alter the conformational ensemble

To probe how the protein dynamics is linked to proton transport within the antiporter module, we next studied how point mutations, that either enhance or impede the proton transport activity in proteoliposomes (*14*), affect the dynamics of the fluorine probe. To this end, we introduced the E132Q substitution (Nqo12^E132Q-N148C-TFA^) at the conserved ion-pair position, which results in an opening of the proton channel and an enhanced proton transport activity across the proteoliposome membranes by electrostatic effects (*14*). Moreover, we studied the effect of the K385I substitution, which impedes the proton transport along the P-side output channel (*14*). We find that the ^19^F-spectrum of the E132Q variant (Fig. 2A) is strongly perturbed by the proton transport-enhancing mutation, resulting in four putative states, **A**-**D**, based on line shape analysis (Fig. S2). The pH-dependent conformations **A** and **B** become apparent at pH<7.5, and titrate with an apparent p*K*_a_ of 7.1 ± 0.2, thus suggesting that the proton affinity of a protonatable cluster increases by >0.7 pH-units relative to the WT construct (Fig. 2E, F). In contrast, the K385I variant (Nqo12^K385I-N148C-TFA^) shows a nearly identical NMR spectrum to that of the WT, suggesting that the point mutation at the P-side exit pathway does not affect the dynamics of the N-side proton input channel (Fig. S3).

### Antiporter conformations depend on the protonation state of central titratable residues

To probe the molecular origin of the protonation-dependent conformational changes, we next performed microsecond MD simulations of the ^19^F-labeled WT and E132Q antiporter constructs embedded in a *n*-dodecyl-β-D-maltoside (DDM) micelle to mimic the conditions in the NMR experiments (see *Method*), together with constant pH-MD (CpH-MD) simulations, which account for protonation changes during the MD simulations. Comparison of the detergent MD simulations with microsecond MD simulations carried out in an *E. coli* cytoplasmic membrane (with POPE:POPG:CDL, see *Methods*, Fig. S4), suggests that the antiporter modules undergo similar hydration changes at functionally relevant regions also in detergents (Fig. S5, Fig. S6).

Interestingly, our CpH-MD simulations indicate that the E132Q substitution drastically shifts the p*K*_a_ of both Lys329 and His241 (Fig. 3E, Fig. S7) along the N-side input channel (*cf*. also Ref. (*18*)), and suggesting that the observed pH-dependence could originate from protonation of these residues. Thus, to probe how the conformational ensemble of the ^19^F-label is linked to the protonation of these residues, we simulated both the protonated (His241^+^) and neutral (His241-δ and His241-ε) tautomeric forms of His241, in combination with the neutral/protonated forms of Lys329 (Fig. 3C, Table S3).

**Figure 3.**
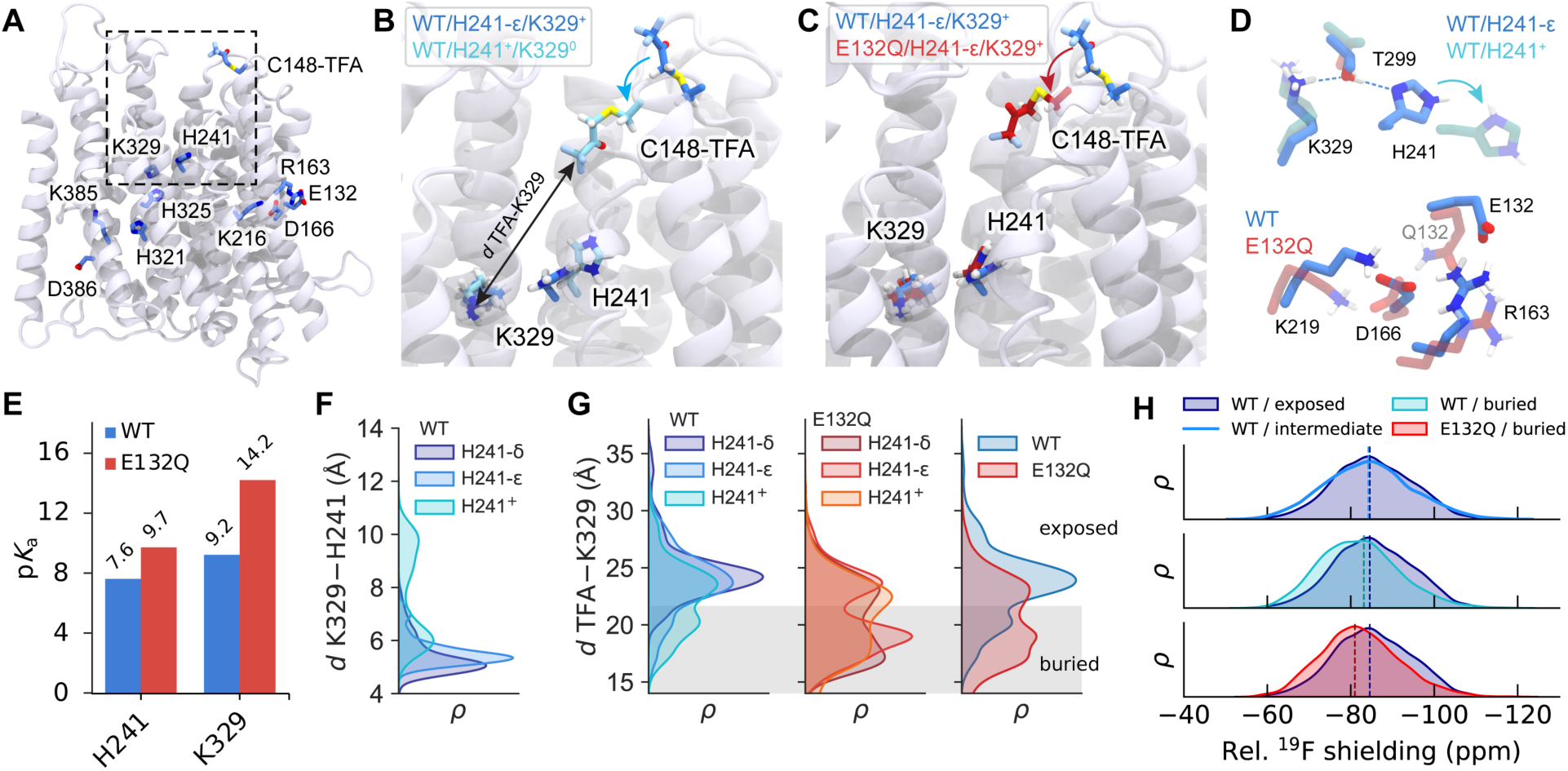
Protonation-dependent dynamics of the NMR-probe in antiporter constructs. (**A**) Structure of Nqo12, showing key residues and the position of the ^19^F-label. (**B**) Conformations of the ^19^F-label in WT-Nqo12 for the H241-ε/K329^+^ (blue, exposed conformation), and H241-p/K329^0^ (light cyan, buried conformation) states. (**C**) Conformations of the ^19^F-label in WT-Nqo12 (blue) and E132Q-Nqo12 (red) in the H241-ε/K329^+^ state. (**D**) Interactions between H241-T299-K329 in different protonation states of H241. (**E**) p*K*_a_ of H241 and K329 in WT-Nqo12 (blue) and E132Q-Nqo12 (red) based on cpH-MD/PBE/MC simulations. (**F**) K329(Cβ)-H241(Cβ) distance distribution for WT-Nqo12. The K329^+^ and K329^0^ states are clustered together for all H241 states. (**G**) Distributions of the ^19^F-label (CF_3_-carbon-K329(Cβ) distance) for WT-Nqo12 (left) and E132Q-Nqo12 (middle) with different protonation states of H241), and combined WT or E132Q in different protonation states (right). (**H**) QM/MM calculations of ^19^F-NMR chemical shifts for exposed and buried ^19^F-label conformations.

Clustering analysis of the MD data suggests that the ^19^F-label can populate 2-3 distinct conformations – an “*inward*-facing” conformation, in which the label points towards the N-side proton channel; an “*outward-facing*” conformation, with a solvent-exposed label pose; as well as a “*partially exposed”* conformation, with the label sampling binding poses between the other two former states (Fig. 3A, B, Fig. S8). Although it is possible that also other conformations appearing on the slower millisecond timescale are involved, our simulations suggest that *inward* and *partially exposed* conformations could result from the pH-and temperature-dependent resonances **A** and **B**, while the temperature-independent broad resonance **C**, could arise from the multiple solvent-exposed conformations of the label.

Our MD simulations further suggest that the His241-ε tautomer forms a hydrogen-bonded network with Lys329 and Thr299 that shifts the histidine towards the N-side proton channel (Fig. 3D, F, Fig. S8), and results in dehydration of the region. These conformations result in a narrow proton input channel, which favors a solvent-facing conformation of the TM5-6 loop, and in turn, leads to an increased population of exposed conformations of the ^19^F-label (Fig. 3, Fig. S8). The ^19^F-label interacts mainly with the TM5-6 loop (residues 145-149), while similar contacts are also observed in the His241-δ form (Fig. S9). However, in stark contrast, protonation of His241 dissociates the His241 – Lys329 – Thr299 contacts (Fig. 3D, F), and drastically increases the hydration level of the channel (Fig. S5). Moreover, the resulting His241^+^ state favors opening of TM5/6-TM11 contacts (Fig. S10), which shifts the TM5-6 loop towards the N-side input channel, and results in a buried ‘*inward’* conformation of the ^19^F-probe (Fig. 3B, G), by contacts to Tyr151, Glu346, and Lys292 (Fig. S9, Fig. S10). In contrast, the His241-δ and H241-ε states show rather high conformational flexibility at the TM5-6 region, with the ^19^F-label adopting a partially shielded conformation. Taken together, these observations further reinforce that the chemical environment sampled by the ^19^F-probe is sensitive to the protonation states of His241 and Lys329.

The E132Q substitution results in an opening of the Glu132-Arg163/Lys216 ion-pair, which increases channel hydration and gives rise to conformational changes of the antiporter around the N-side input channel (Fig. S5, Fig. S10). The channel hydration and conformational dynamics around the TM7a and TM5-6 region in the E132Q variant are similar to the His241^+^ state of the WT-Nqo12, thus indicative of a putative conductive channel state. Overall, the conformational and hydration changes result in a more buried, *inward* conformation of the ^19^F-probe (Fig. 3G, Fig. S10), as also observed in in the His241^+^ state of the WT. Indeed, principal component analysis (PCA) further supports that the ^19^F-probe in the E132Q variant result in conformations that overlap with the H241^+^ state of the WT, but are distinctly different from other tautomeric forms of His241 in the WT (Fig. S11, Fig. S12, Fig. S13). Moreover, our contact analysis reveals multiple interactions of the ^19^F-label around the N-side input channel in all microstates of His241/Lys329 in E132Q-Nqo12 (Fig. S9, Fig. S13). In contrast, the His241-δ/His241-ε states of WT-Nqo12, form interactions mainly with the TM5-6 loop (Fig. S9, Fig. S12). Taken together, these interactions could result from a buried conformation of the ^19^F-probe, and possibly result in additional NMR resonance, as experimentally observed for the E132Q-Nqo12 variant (Fig. 2F).

To further correlate the ^19^F-NMR spectra with conformational states of Nqo12, we simulated the ^19^F-NMR spectra for the three main conformations of the NMR label observed in our MD simulations (*inward*-, *intermediate*- and *outward*-facing) using hybrid quantum/classical molecular dynamics/molecular dynamics (QM/MM-MD) calculations, initiated from our microsecond MD simulations. Although the quantitative *ab initio* prediction of NMR chemical shifts is highly challenging (*32*, *33*), we note that our QM/MM-MD simulations of the distinct ^19^F-label conformations show a systematic upfield shift of the computed ^19^F chemical shifts in the *outward* label position, as compared to the *inward* position at the KT2/pcSseg-2/MM level (*34–36*) (see *Methods*, Fig. 3H), leading to a distinct difference (+3.5 ppm) in chemical shifts between the WT-Nqo12 and E132Q-Nqo12, in qualitative agreement with the experimentally observed spectral changes (Fig. 2A). These observations, together with our classical molecular simulations, further support our assignment of the label poses and experimental observations.

### Functional mutations increase proton transport and remove pH-dependence of the proton uptake site

To experimentally validate the identified proton uptake site, as revealed by our multiscale simulations, we introduced K329M and H241A substitutions into the antiporter-like module, followed by expression and purification of the constructs. To quantify the proton conduction, we employed proteoliposome experiments, in which we co-reconstituted these variants with ATP synthase (F_o_F_1_-ATPase) to generate a PMF upon addition of ATP, and monitored pH-changes optically with the pH-sensitive ratiometric dye pyranine (HPTS) by fluorescence spectroscopy (Fig. 4E). We find that both the K329M and H241A substitutions result in drastically increased proton conduction rates across the proteoliposome membrane relative to the WT-Nqo12 (Fig. 4E-G), indicative of disruption of a local proton gate. Moreover, the K329M and H241A variants completely abolish the pH dependence of the **A** / **B** populations in NMR experiments (Fig. 4A-D), further suggesting that the identified His241/Lys329 cluster indeed forms a key proton uptake site that controls proton conduction across the antiporter module.

**Figure 4.**
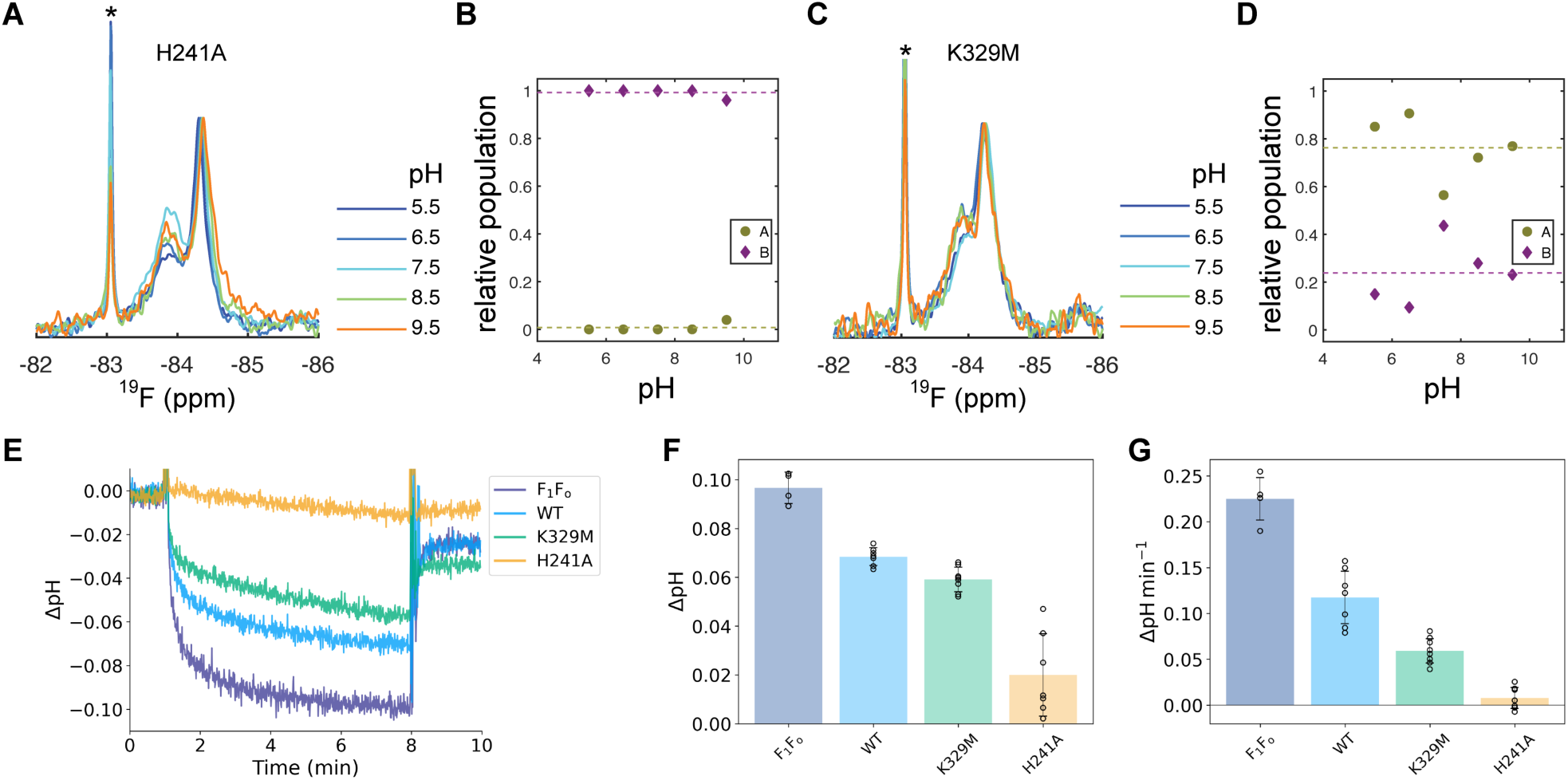
Functional residues involved in the proton uptake. **(A,C)** ^19^F-spectra and (**B,D**) pH-dependence of the **A**/**B** ratio for the (**A,B**) H241A and (**C,D**) K329M-Nqo12 variants. (**E**) Proteoliposome assays, monitoring the pyranine signal (acidification) in the inner lumen of the vesicles containing ATP synthase (F_1_F_o_) alone as control, or co-reconstituted with Nqo12-WT or the variants. The assay is started after 1 min by addition of ATP (see *Methods*). (**F**) The ΔpH, fitted from data in (**E**), using a single exponential function. (**G**) Initial rates, obtained from linear fitting of the first 20 seconds of data in (**E**).

## DISCUSSION

We have identified here a key proton uptake site in the terminal antiporter module Nqo12 (NuoL/ND5) that undergoes conformational changes upon proton transport. Based on combined NMR experiments, mutagenesis experiments, and multiscale molecular simulations, we showed that protonation of the identified His241/Lys329 site is regulated by opening of a key conserved ion-pair element, previously linked to proton transport in Complex I (Fig. 5A) (*7*, *18*, *19*). Our current findings suggest that the protonation of this cluster regulates the proton conduction by hydration (wetting) and dehydration (dewetting) of the N-side input channel. The site could thus form an important gating element during the redox-driven proton pumping process of Complex I, in which proton leaks must be prevented by blocking proton transport in the thermodynamically favored directions (P-side → N-side at high PMF).

**Figure 5.**
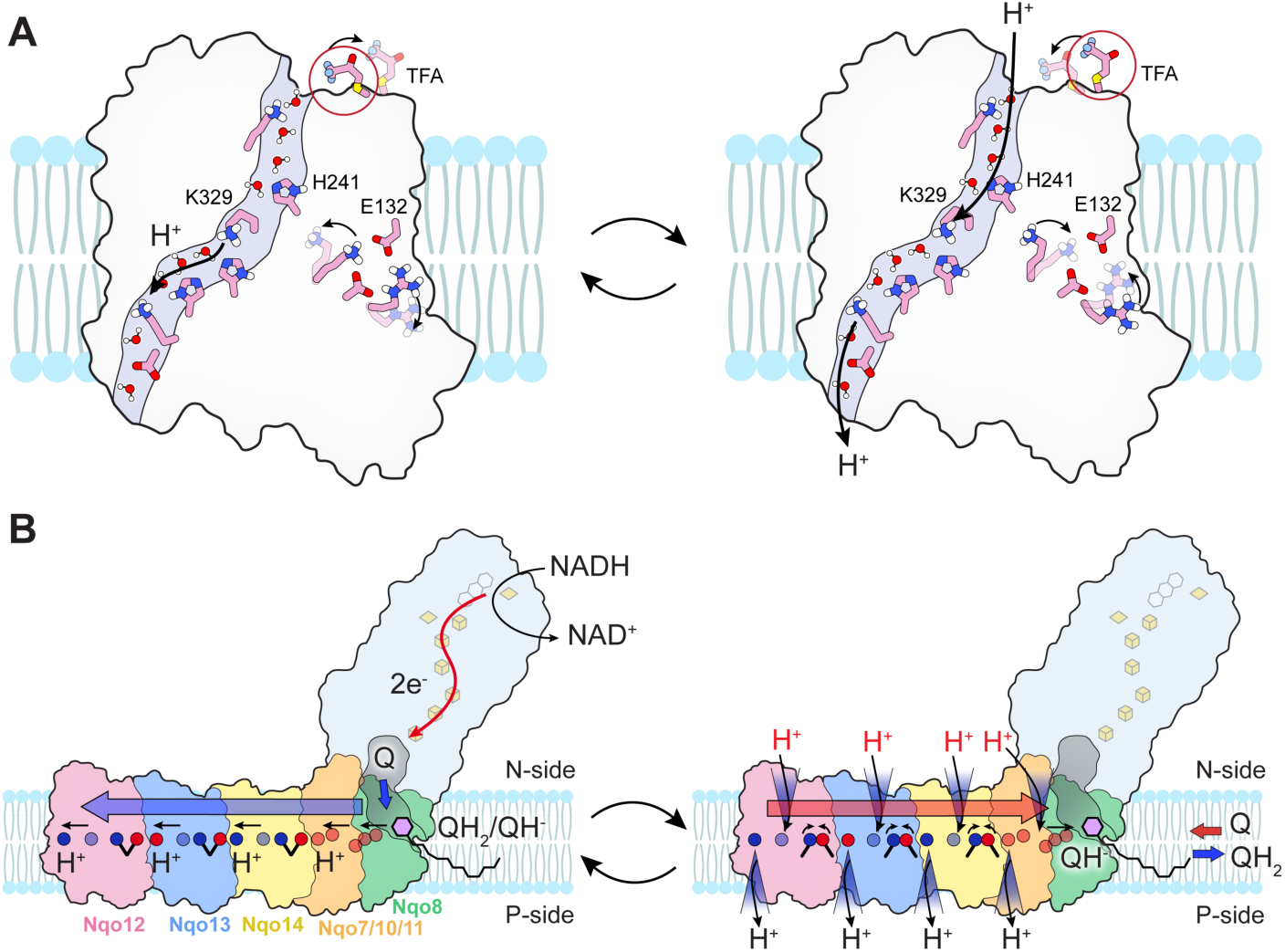
Schematic model of putative conformational gating transitions underlying proton transport in antiporter-like modules and in Complex I. **(A)** Opening of ion-pair element favors lateral proton transfer within the antiporter module. Protonation change of the His/Lys cluster shifts the position of the NMR probe to the buried conformation. Proton uptake to the His/Lys cluster couples to proton release across the P-side, favoring ion-pair closure, while shifting the position of the NMR probe to the exposed conformation. Consecutive transitions between ‘frustrated’ open ion-pair / deprotonated Lys/His cluster and closed ion-pair / protonated Lys/His cluster could drive the proton flux through the antiporter module, upon applying a PMF. **(B)** *Left:* Q-reduction and Q-diffusion induces stepwise propagation of a protonation wave across Nqo7/10/11/8, Nqo14, Nqo13, and Nqo12, and result in sequential ion-pair opening and lateral proton transfer. *Right*: The shifted protonation states along the lateral hydrophilic axis favor opening of proton gates along the N- and P-side pathways, followed by stepwise closing of the ion-pair. The process back-propagates from Nqo12, *via* Nqo13, Nqo14, and Nqo7/10/11/8 to the membrane bound quinol site. Upon re-protonation of a putative QH^-^ species, QH_2_ is released to the membrane, and Complex I returns to the resting state.

Our NMR experiments together with molecular simulations revealed that ion-pair opening results in a >0.7 p*K*_a_ shift of the identified His/Lys-cluster. However, the activated channel, studied here *via* the E132Q substitution, also resolved a fourth resonance **D,** in addition to the broad temperature-insensitive state **C**, and the temperature- and pH-dependent conformations **A** and **B**. While we did not investigate the nature of this conformation here in detail, we suggest that state **D** could arise from the perturbed protonation of Lys329 (Lys329-NH_2_^0^ / Lys329-NH_3_^+^) in addition to the tautomeric states of His241 (H241-ε, H241-δ, His241^+^), which are favored by the E132Q substitution, and could populate a mixture of open and closed channel conformations of TM7a/b and TM8.

Our findings provide molecular insight into how protonation dynamics are linked to distinct protein conformational changes in TM helices that regulate proton transport in Complex I. While difficult to resolve in cryo-EM or X-ray experiments, subtle conformational changes at and around the broken helices TM7a/b and TM12a/b were suggested based on molecular simulations (*16*, *18*, *20*, *21*, *37*) to control the proton uptake during the pumping process. To this end, Di Luca *et al.* (*18*) suggested that the proton uptake by His241 and Lys329 is controlled by a steric bottleneck around Phe333/Leu373 in Nqo12 (*T. thermophilus* Complex I numbering), leading to conformational changes along the N-side input channel (*18*). Röpke *et al.* (*20*) further observed subtle conformational changes in TM7a/b and TM12a/b helices upon protonation changes along the lateral proton channel, whereas Badolato *et al.* (*15*) showed that mutation of the non-polar Leu/Phe residues controlled the channel hydration and rate of transmembrane proton transport in proteoliposomes. Together with work by Beghiah *et al.* (*14*), these findings provide support that the ion-pair elements control the proton transport in the antiporter modules of Complex I. Moreover, Hoeser *et al.* (*21*) recently observed conformational changes in His241 (His254^NuoL^ in *E. coli* Complex I) upon introduction of a disease-related mutation (F111L) close to the ion-pair element, further supporting its functional relevance, whereas Saura *et al.* (*16*) showed that broken helix elements TM7a/b and TM12a/b may undergo a conformational breathing motion that could regulate proton accessibility to the central channel, but possibly also short-circuiting protonation to the ion-pair element upon harmful mutations.

The importance of the analogous His241 residue was also recognized in the histidine switch model, initially suggested based on cryo-EM and MD simulations of the Mrp transporters (*38*) and later expanded to respiratory Complex I (*22*) (*cf*. also Ref. (*23*)). This model suggested that the residue analogous to His241 could, by its tautomeric protonation state, specifically H241-ε or H241-δ, regulate proton uptake via the central proton channel or, both to and from, the ion-pair element to the N-side (*22*, *23*). While our model also supports that this histidine residue undergoes protonation-dependent conformational dynamics, our data also reveal that the nearby lysine (Lys329) is tightly coupled to this protonation process, and *vice versa*. Moreover, the proposed bidirectional proton transfer to and from the ion-pair element to the N-side could create a short-circuit pathway stimulating proton leaks that impede the overall pumping efficiency, particularly upon introduction of disease mutations near the ion-pair elements (*16*).

We suggest that the identified functional elements resemble a field-effect transistor, in which a tunable electric field controls the current flow across the gate. By analogy, the ion-pair element modulates the electric field at the identified protonation site, controlling its protonation state. In turn, this creates an electric field between the N-side bulk and the interior of the antiporter module, which controls wetting/dewetting transitions of the channel (by water dipole / electric field interactions), and thus the proton current across the gate.

According to the electrical wave propagation mechanism (*7*, *14*, *18*, *19*), sequential opening of the conserved ion-pair elements results in the propagation of a lateral protonation wave along the proton hydrophilic axis by electrostatic effects (Fig. 5B). Based on our combined data, we find that opening of the conserved ion-pair element electrostatically upshifts the p*K*_a_ of the central His241/Lys329 cluster, which in turn triggers proton uptake along the hydrated N-side input channel and lateral water-mediated proton transfer via His241/Lys329 to the terminal Lys385/Asp386. Proton release across the P-side exit channel of TM12a/b could stimulate further proton uptake along the N-side input channel, which in turn shifts the protonation state of the His241/Lys329 cluster, leading to an electrostatic repulsion that closes the ion-pair element. In turn, this process could initiate the back-propagation of similar proton release, uptake, and ion-pair closures in Nqo13 and Nqo14, propagating all the way to Nqo7/8/10/11, and leading to the release of the membrane-bound quinol to the membrane (Fig. 5B). Although the exact proton output sites still remain debated, all models support that Nqo12 (NuoL/ND5) in the canonical Complex I superfamily is involved in both proton uptake and release (but see below).

The importance of conformational switching of His241 was also recently supported by structural studies of the *Bos taurus* Complex I (*12*), suggesting that the residue might form a ‘master-switch’ that controls the proton pumping. However, due to the lack of an analogous conserved histidine residue in ND2 (Nqo14), it was proposed that ND2 (Nqo14) cannot be involved in proton uptake, thus partially contrasting earlier findings by Grba *et al.* (*28*), in which water-mediated contacts leading from the N-side were observed. Based on their findings, Fischer *et al.* thus favored a model in which all protons are pumped via ND5, with proton uptake taking place in ND4 and ND5 (*cf.* also (*39*)), following the proposal by Sazanov and co-workers (*8*, *9*). Importantly, however, water contacts, or lack thereof, cannot directly be used to conclude the feasibility of proton transport – with both aquaporin, where the water channel is always present, and cytochrome *c* oxidase, where no experimentally observed pathway connecting to the active sites has been observed, serving as counter examples. Nevertheless, while likely mediating “a protonation signal”, *e.g.,* as suggested by the wave propagation mechanism, future work needs to probe the molecular details of ND2 (Nqo14) in proton uptake and release.

Efficient redox-driven proton pumping in Complex I is based on tightly coupled energy transducing steps, in which small redox and p*K*_a_ gaps between individual microstates secure microscopic reversibility, and minimize dissipation effects. Thermodynamically, the proton pumping in Complex I is expected to evenly distribute the *ca*. 0.8 V redox free energy (*E*_m_(NADH)=-320 mV → *E*_m_(Q)=+90 mV) from the NADH-driven Q-reduction within the 200 Å wide membrane domain. The electrical wave mechanism introduces microscopic reversibility by creating tight coupling within and between each antiporter-like subunit, thus limiting each local coupling effect to modest (5-10 kcal mol^-1^) energetic effects, required to overcome the electrochemical work of transferring the proton against a 200 mV (4.6 kcal) PMF. However, models in which all four protons are pumped by the terminal Nqo12 (NuoL/ND5) subunit may require that the full redox energy (>19 kcal mol^-1^) is transduced in a single step to the terminal antiporter module, and could thus be more prone to dissipation effects and proton leaks. As Complex I does not accumulate one-electron reduced semiquinone intermediates, it remains difficult to rationalize how a single redox stroke could result in ejection of four consecutive proton transfer steps, even if features of the wave propagation model (*7*, *14*, *18*, *19*) are considered. Importantly, the “ND5-only”–like models are also inconsistent with the experiments by Dröse *et al.* (*17*), showing that the ND4-ND5 knockout of Complex I is competent in catalyzing a redox-driven proton pumping with a reduced stoichiometry of approximately 1 H^+^/e^-^.

Complex I has a modular structure, with large variations in both electron input sites, with ferredoxin-dependent variants and NADH-dependent variants mediating electron transfer both with and without the N-module, together with variations in the structure of the antiporter-like subunits (*40*). For example, in the photosynthetic NDH-1MS (*41*) the proton pumping Nqo12 (NuoL/ND5) module is replaced with a distinct NdhF3-CupA/S subunit, which lacks key functional elements of the proton channel, while the protein complex still pumps protons and generates a PMF (*42*). Also, variants with only a single proton channel (Ech) or with additional proton channels (*Thaumarchaeal 2M* clade) exist (*40*, *43*). Together with different redox gaps between the electron donors and acceptors, such Complex I homologs could give rise to different redox-driven proton pumping stoichiometries within the superfamily. More generally, this implies that while all Complex I isoforms could employ a generic mechanism of a protonation wave across the membrane domain, the individual isoforms might use distinct principles to relax the “proton-loaded” states. Indeed, similar functional variations are also observed in other bioenergetic machines, such as the superfamily of heme-copper oxidases (HCOs) (*44*, *45*), which employ one or two proton input channels, despite all isoforms relying on variations of a water-mediated proton bifurcation mechanism to pump protons (*cf*. Refs. (*46–49*) and refs. therein). Future work will need to identify the generic conserved elements required for long-range redox driven proton pumping in Complex I by integration of biochemical, structural, spectroscopic, and computational experiments.

In summary, we have combined here NMR spectroscopy and site-directed mutagenesis experiments with multiscale molecular simulations to resolve how protonation changes in the terminal antiporter-like module (Nqo12/NuoL/ND5) of Complex I are linked to protein conformational dynamics. We identified a conserved histidine/lysine cluster on the broken helix element TM7a and TM8 (His241/Lys329^Nqo12^) that undergoes protonation changes, controlling the conformational state of surrounding helices, and regulating proton transport across the subunit. We showed that the apparent p*K*_a_ of the identified cluster shifts from 6.4 to 7.1 upon opening of a conserved ion-pair element in the antiporter module, while mutation of either the histidine or lysine residues resulted in a complete loss of the pH-dependent conformational changes. The mutated antiporter module also showed increased conductivity in our proteoliposome assays, in agreement with the role of the histidine and lysine residues in the proton transfer regulation. Our combined findings are consistent with key features of the electrical wave propagation mechanism (*7*), in which ion-pair dynamics and proton uptake through the broken helix elements result in propagation of a protonation wave across the antiporter module. The identified histidine/lysine cluster is thus likely to function as an important regulatory switch in this process. Taken together, our findings highlight key molecular locks and gates that control the long-range redox-driven proton pumping process in Complex I. The identified elements show interesting similarities to electrical gates, suggesting that the antiporter module functions similarly to a molecular field-effect transistor, regulating transmembrane proton currents.

## MATERIALS AND METHODS

### Molecular dynamics simulations

Classical molecular dynamics (MD) simulations of the ^19^F-TFA labelled WT-Nqo12 (Nqo12^N148C-TFA^) and E132Q-Nqo12 (Nqo12^E132Q-N148C-TFA^) variants were performed, using a model constructed based on an engineered Nqo12 subunit of *T. thermophilus* Complex I (PDB ID: 4HEA (*2*)), following removal of the transverse helix (residues 516-606). To mimic the NMR experiments, the protein was embedded in an *n*-dodecyl-β-D-maltoside (DDM) detergent micelle (*N*=250 detergent molecules), solvated with TIP3P water molecules and 150 mM NaCl, as guided by CHARMM-GUI(*50*), and resulting in MD systems of 250,000 atoms (Fig. S4A). The simulations were compared to simulations of the same systems carried out in a POPE:POPG:CDL (7:2:1 ratio) lipid, with around 125,000 atoms (Fig. S4B). The protein, detergent and solvent molecules were described at the CHARMM36 force field level (*51*), whereas the ^19^F-TFA label was parametrized based on DFT calculations (see below). MD simulations were performed at 1 bar pressure at 310 K, using an *NPT* ensemble, with a 2 fs integration timestep, and treating long-range electrostatic interactions by the Particle Mesh Ewald method. For each construct, different protonation states of residues H241/K329/K385 were considered. 1 μs MD simulations of each construct and condition were performed in duplicates (see Table S3). All classical MD simulations were performed with NAMD 3.0 (*52*), and DFT calculations were performed with TURBOMOLE v7.7 (*53*). Visualization was carried out with VMD (*54*) and UCSF ChimeraX (*55*), and data was analyzed with VMD (*54*) and MDAnalysis (*56*). Clustering analysis was performed to examine the conformations of the ^19^F-TFA label in the different constructs and protonation states. To remove contributions from overall protein motion, the protein backbone was first aligned, and the probe conformations were clustered using the quality threshold algorithm, as implemented in VMD (*54*), using an RMSD cutoff of 2.5 Å, based on the ^19^F-TFA-Cys148 heavy atoms. The simulation data were analyzed for the clustering analysis with a frequency of 1 frame per ns, with most populated clusters selected for the subsequent QM/MM simulations (see below). See Table S4 for clustering analysis.

### Force field parametrization of ^19^F-TFA label

Force field parameters for the ^19^F-TFA label were derived based on DFT calculations. To this end, a model system, comprising Cys-CH_3_-C=O-CF_3_, was optimized at the B3LYP-D3/def2-SVP level (*57–60*). Following structure optimization, atomic partial charges were calculated at the B3LYP-D3/def2-TZVP level using the restrained electrostatic potential (RESP) (*61*) method, while Lennard-Jones and bonded parameters were derived from the CHARMM36m force field (*51*).

### Constant pH-MD simulations

Constant-pH MD (cpH-MD) simulations were performed based on MD simulations of Nqo12 in a POPE:POPG:CDL (7:2:1 ratio) lipid membrane by adapting data and methodology from previous simulations (*14*). In this regard, 16 cpH-MD replicas were performed in a pH range 2.8-12.1 based on pre-equilibrated MD structures. Each simulation comprised 1000 minimization steps, followed by 5000 cycles of production steps (5000 steps each), and 7500 non-equilibrium switching steps, leading to a *ca*. 400 ns sampling. Initially, all residues were sampled in their standard reference state, and by enhancing protonation changes of H241, K292, and K329 using an external protonation bias of 10 kcal mol^-1^. Based on the cpH-MD sampling, 1500 structures from different pH conditions were selected for computation of p*K*_a_ values using Poisson-Boltzmann (PB) electrostatic calculations coupled to Monte Carlo (MC) sampling. In the initial calculation, all titratable groups were modeled, followed by a focused p*K*_a_ calculation of residues along the proton pathway (H341, K147, H88, H241, K216, D166, E132, K292, H321, H325, K329, E490, K385, D386).

A final MC sampling was performed with all conformations re-weighed by their respective protonation probability and electrostatic energy. The cpH-MD simulations were performed as implemented in NAMD (*62–64*), while the p*K*_a_ computations were performed using an in-house version of APBS/Karlsberg+ (*65–68*).

### Hybrid QM/MM calculations

Hybrid quantum/classical (QM/MM) simulations were performed to calculate chemical shifts of the ^19^F-TFA probe in the different constructs. To this end, classically relaxed snapshots were selected from the MD simulations (Table S3), based on the clustering analysis. The QM/MM systems were truncated to around 65,000 atoms, with the QM region comprising 220-270 atoms depending on the conformation of the NMR probe, which contained the protein residues within the first sphere of the ^19^F-TFA label, together with water molecules around 7 Å of the probe (Table S5). The QM/MM partition was described by a link-atom approach, with link atoms positioned between the Cα and Cβ atoms of protein residues (and between Cα-C and Cα-N for backbone atoms), using an additive electrostatic embedding scheme. The system was initially minimized for 50 steps using the Adapted-Basis Newton-Raphson method, followed by 5 ps of QM/MM MD at *T*=310 K, with a 1 fs integration timestep. All atoms within 10 Å of the QM region were allowed to relax, while keeping the rest of the system fixed. Initial minimization and QM/MM-MD sampling were performed at the B3LYP-D3/def2-SVP level (*57–60*), with classical point charges described by the CHARMM36 force field (*51*). Chemical shifts were calculated at the KT2/(all)pcSseg-1(all)/pcSseg-2(CF_3_) level (*34*, *35*), based on conformations extracted from the QM/MM simulations (see Fig. S14). Computed absolute ^19^F-shieldings were shifted to the average experimental chemical shifts of resonance **A** of the ^19^F-TFA in the WT-Nqo12 spectrum. All QM/MM calculations were performed with CHARMM c38b1 (*69*)/TURBOMOLE 7.7 (*53*), coupled by a Python interface (*70*).

### Cloning, expression, ^19^F-labeling and purification

Following our previously established approach (*14*), the truncated Nqo12 variants (with the C-terminal amphipathic helix, residues 516-606, removed) were expressed. The codon-optimized truncated *nqo12* gene from *T. thermophilus* was incorporated into a pWALDO vector (*71*) carrying an ampicillin resistance cassette and the construct was expressed with a C-terminal linker (GSAGS), connecting to a TEV-cleavage site, followed by superfolder green fluorescent protein (sfGFP (*72*)), and a His_10_-tag. Substitutions were incorporated into pWALDO-*nqo12* employing a QuikChange procedure. Designed primers for the constructs are listed in Table S1. The Nqo12 constructs were expressed and purified as described previously (*14*). Briefly, the plasmid was transformed into *E. coli* Lemo21 competent cells (*73*), incubated in LB overnight with appropriate antibiotics. On the next day the culture was diluted to an OD_600_ of 0.05 into TB medium with 0.1 mM L-rhamnose, grown at 37°C to an OD_600_ of 0.5, induced with isopropyl β-D-1-thiogalatopyranoside (IPTG), with protein expressed overnight at 25°C. Cells were harvested on the next day and the pellet was stored at −80°C until purification.

For protein purification (Fig. S15), cells were resuspended in buffer A (25 mM HEPES pH 7.5, 150 mM NaCl, see Table S6) supplemented with 0.1 mg ml^-1^ AEBSF protease inhibitor and 10 µg ml^-1^ DNase I. Cells were then lysed by passing them three times through a high-pressure homogenizer at ∼15 000 psi. Cell debris was removed by high-speed centrifugation at 25,000 g for 25 min and membranes were collected by ultracentrifugation at 165,000 g for 2 h. Membranes were resuspended into buffer A at a concentration of ∼6 mg ml^-1^ and solubilized with 0.8% *n*-dodecyl-β-D-maltoside (DDM) or with 2% Lauryl Maltose Neopentyl Glycol (LMNG) for proteoliposomes experiments, and incubated for 1 h at 4°C followed by ultracentrifugation at 165,000 g for 30 min to remove undissolved membranes. The protein was then applied to a Ni-NTA column using an FPLC, and the supernatant was passed over the column 5 times, to increase the amount of bound protein. The column was washed with buffer B (25 mM HEPES pH 7.5, 300 mM NaCl, 0.05% DDM, Table S6), supplemented with increasing amounts of imidazole using a step gradient (20 mM, 75 mM, 125 mM and 150 mM imidazole concentration, each wash was conducted with at least 10 column volumes). The protein was eluted in buffer C (buffer B supplemented with 300 mM imidazole), concentrated to 2.5 ml using a 50 kDa MWCO concentrator tube and buffer was exchanged to SEC buffer (buffer D: 25 mM HEPES pH 7.5, 300 mM NaCl, 0.05% DDM) using a PD10 desalting column. 1 mg of TEV protease was added and cleavage proceeded overnight at 4°C. On the next day, TEV protease and the cleaved sfGFP-His-tag were removed from the sample using reverse IMAC and the protein was concentrated to ∼1 ml.

Samples were ^19^F-labeled prior to size exclusion chromatography (SEC), following a previously published protocol (*24*). To 1 ml of sample, with a typical protein concentration of 10 – 50 µM, 1 mM of dithiothreitol (DTT) was added, and the sample was incubated at 4°C for 15 min. Next, 50 mM 3-bromo-1,1,1-trifluoroacetone (BTFA) was added and the sample was further incubated for 1 h at 4°C. Excess label was removed in the final SEC purification step. However, we noted that a fraction of free label was retained even after SEC purification. Based on the observed ^19^F chemical shift, which is downshifted relative to BTFA in water (*74*), we suspect that free label partitions to some degree into DDM micelles and is thus carried over to the final sample. Finally, the sample was purified by SEC using a HiLoad 16/600 Superdex 200 pg column, the peak fraction was collected and concentrated to ∼200 µl. Preparations for proteoliposome experiments excluded sfGFP cleavage and ^19^F-labelling, unless stated otherwise.

To exclude possible spurious ^19^F-labeling, we also conducted the labeling protocol for the Nqo12^H241C^ variant, for which the buried residue H241 is not accessible to BTFA. Indeed no ^19^F-resonances were observed for this construct (Fig. S1B), showing that the approach does not produce spuriously labeled sites.

### NMR sample preparation

To exchange buffer, ∼200 µl sample was diluted into 15 ml of the appropriate buffer, concentrated back to ∼200 µl, and the procedure was repeated once. 10% D_2_O was added to the sample for frequency locking and the NMR sample was placed in a 3 mm tube. The sample conditions were 35 – 100 µM protein, 25 mM buffer (MES for pH 5.5 and 6.5, HEPES for pH 7.5, and Tris/HCl for pH 8.5 and 9.5), 300 mM NaCl, 0.05% DDM, and 10% D_2_O.

### NMR experiments

NMR experiments were conducted on a Bruker Avance Neo 500 MHz (471 MHz ^19^F frequency) equipped with a nitrogen-cooled triple-resonance TCI probe head, for which the ^1^H channel could be tuned to ^19^F frequency, and a Bruker Avance Neo 600 MHz (564 MHz ^19^F frequency) spectrometer equipped with a helium-cooled quadruple-resonance QCI-P probe head. Experiments were conducted with a sweep width of 20 ppm, an acquisition time of 106 ms and a recycling delay of 1.5 s at a sample temperature of 313 K, if not indicated otherwise. Typically, for 1D experiments 16,000 transients were collected, while 7000 transients were collected for the CEST experiment. Raw data were processed using NMRPipe/NMRDraw (*75*).

### CEST experiment

A CEST experiment was acquired for a B_1_ field of ν_1_ = 10 Hz and 33 offset frequencies [-10000, −200, −175, −150, −125, −100, −75, −50, −25, 0, 25, 50, 75, 100, 125, 150, 175, 200, 225, 250, 275, 300, 325, 350, 375, 400, 425, 450, 475, 500, 525, 550, 10000] Hz, where 0 Hz was set to the chemical shift of the most intense peak. Peaks were integrated using an in-house Matlab script and integrals were referenced to the average value obtained for ±10,000 Hz. CEST profiles for resonance **C** and **A**/**B** were simultaneously fitted to the data, employing a two-state exchange model using numerical integration of the Bloch-McConnell equation (*76*, *77*) (Eq. 1) with global parameters to model population of conformation **C**, as *p*_C_ (= 1 – p_A/B_), the exchange rate, *k*_ex_ = *k*_C,A/B_ + *k*_A/B,C_, for which *p*_A/B_ = *k*_C,A/B_ / *k*_ex_ and *p*_C_ = *k*_A/B,C_ / *k*_ex_, and the chemical shift difference between conformation **C** and **A**/**B**, Δν. The relaxation rate *R*_1,C_ was assumed to be equal to *R*_1,A/B_, while *R*_2,C_ and *R*_2,A/B_ were treated as distinct parameters based on,

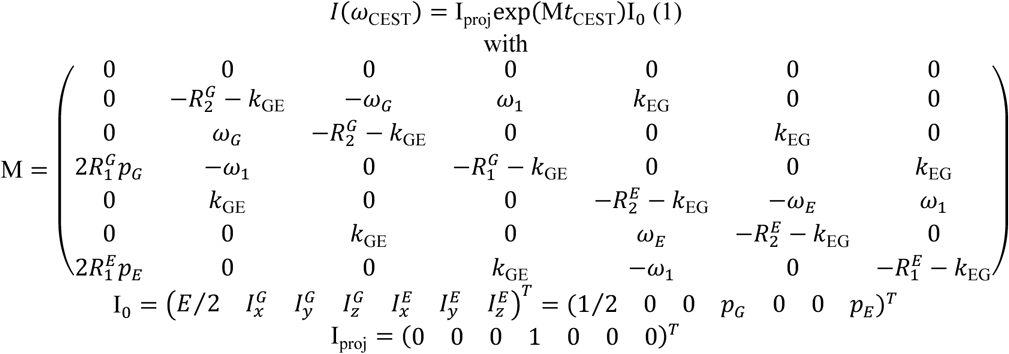

for which **I_0_** is the initial magnetization of the observed ground (G) and excited (E) state and **I_proj_** extracts the observable z-component from the final modulated magnetization. ω_G/E_ = 2πB_0_ (δ/B_0_ – δ_G/E_) is the offset of state G or E, with resonances at δ_G/E_ (in ppm) from the CEST irradiation frequency, and the intensity ω_1_ = 2πν_1_ at δ (in Hz) and Δν = δ_E_ – δ_G_. Note that for a fit of state **C** CEST profile, C ≘ G, and A/B ≘ E in Eq. 1 and *vice versa* when the **A**/**B** state CEST profile was fitted. Measurement uncertainties for the CEST parameters were obtained by a Monte Carlo algorithm, in which 1000 artificial CEST profiles were generated within the errors of the experimental dataset. The uncertainty of the CEST parameters was determined from the standard deviation of the parameters obtained from fitting Eq. 1 to the artificial datasets.

### ^19^F spectral analysis

To obtain populations of the conformations probed by the ^19^F-label, spectra were referenced to the BTFA peak, and four (for Nqo12-E132Q) or three (for other constructs) Lorentzian line shapes were fitted to the data, where we note that fits with fewer Lorentzians could not represent the experimental data. Line shapes were fitted globally to the spectra acquired at different pH utilizing the fminsearch routine in Matlab (version 2022b) employing in-house scripts. We assumed constant line width for each line shape under all conditions, but allowed for variability in the peak position and integral.

To obtain an apparent p*K*_a_ of the constructs, we plotted the two pH-sensitive populations obtained from the integral of the Lorentzians against the pH and fitted a sigmoidal function to the data, for which the position of the inflection point corresponds to the apparent p*K*_a_. Measurement uncertainties for the populations obtained from the line shape fits were determined by bootstrap analysis, and errors for the sigmoidal fit were obtained from a Monte Carlo simulation of 1000 artificial datasets generated within the standard deviation of the populations.

### Proteoliposomes assays

Liposomes were prepared from soybean Type II-S lipids, and resuspended in the proteoliposomes buffer (2 mM HEPES, 50 mM KCl, 5 mM MgCl_2_, pH 7.3, see Table S6). Multilamellar formations were removed by 7 freeze/thaw cycles, while pyranine (8-hydroxypyrene-1,3,6-trisulfonic acid, HPTS), was incorporated in the inner lumen of the vesicles. Liposome extrusion was performed 21 times to select for 100 nm of vesicle diameter, using pored membranes (Nuclepore membranes, *Whatman Ltd*). The protein reconstitution was performed with the LMNG-autoinsertion reintegration (LAiR) approach (*78*), by adding LMNG-purified *E. coli* F_1_F_o_-ATP synthase, expressed and purified using a previously established protocol (*15*), and LMNG-purified Nqo12-WT or variants, in a ratio of 3:4 (F_1_F_o_-ATP synthases: Nqo12) protein molecules per vesicle. The sample was incubated at room temperature for 30 min before external HPTS was removed using a PD-10 desalting column, and ultracentrifugation for 30 min at 50,000 rpm at 4°C. The proteoliposome pellet was resuspended in the proteoliposome buffer at a lipid concentration of 20 mg ml^-1^. Measurements were performed using a Cary Eclipse Fluorescence Spectrophotometer (Agilent Technologies). HPTS excitation wavelengths were set at 405 nm and 460 nm, while emission was followed at 510 nm at 37°C. Plastic cuvettes were used in a final volume of 500 µl of proteoliposome buffer, with 5 µl of proteoliposome resuspension, and 1 nM valinomycin. After one minute of equilibration, the assay was initiated by addition of 0.04 mM ATP, and the generated ΔpH was dissipated at minute 7 by addition of gramicidin, with the HPTS signal converted to pH changes, based on previous calibration (*15*).

## Acknowledgments

We thank the Ulrika Brath and the Swedish NMR Centre at the University of Gothenburg, and Prof. Remco Sprangers, University of Regensburg, for providing access to NMR spectrometers, and Adel Beghiah for insightful discussion and technical help during early stages of the project.

## Author contributions

Conceptualization: VRIK; Methodology: JL, PS, VRIK; Investigation: JL, PS, SB, APGH; Formal Analysis: JL, PS, SB, APGH; Visualization: JL, PS; Funding acquisition: VRIK; Project administration: VRIK; Supervision: VRIK; Writing – original draft: JL, PS, VRIK; Writing – review & editing: JL, PS, SB, VRIK.

## Data, code, and materials availability

All data are available in the manuscript or the supplementary materials.

## REFERENCES

1. V. R. I. Kaila, M. Wikström, Architecture of bacterial respiratory chains. Nat Rev Microbiol 19, 319–330 (2021).

2. R. Baradaran, J. M. Berrisford, G. S. Minhas, L. A. Sazanov, Crystal structure of the entire respiratory complex I. Nature 494, 443–8 (2013).

3. M. Verkhovskaya, N. Belevich, L. Euro, M. Wikstrom, M. Verkhovsky, Real-time electron transfer in respiratory complex I. Proc. Natl. Acad. Sci. U. S. A. 105, 3763–3767 (2008).

4. L. Euro, G. Belevich, M. Verkhovsky, M. Wikstrom, M. Verkhovskaya, Conserved lysine residues of the membrane subunit NuoM are involved in energy conversion by the proton-pumping NADH: ubiquinone oxidoreductase (Complex I). Biochim. Biophys. Acta-Bioenerg. 1777, 1166–1172 (2008).

5. M.-C. Kao, E. Nakamaru-Ogiso, A. Matsuno-Yagi, T. Yagi, Characterization of the Membrane Domain Subunit NuoK (ND4L) of the NADH-Quinone Oxidoreductase from*Escherichia coli*,. Biochemistry 44, 9545– 9554 (2005).

6. J. Michel, J. DeLeon-Rangel, S. Zhu, K. Van Ree, S. B. Vik, Mutagenesis of the L, M, and N Subunits of Complex I from Escherichia coli Indicates a Common Role in Function. PLoS ONE 6, e17420 (2011).

7. V. R. I. Kaila, Long-range proton-coupled electron transfer in biological energy conversion: towards mechanistic understanding of respiratory complex I. J. R. Soc. Interface 15, 20170916 (2017).

8. V. Kravchuk, O. Petrova, D. Kampjut, A. Wojciechowska-Bason, Z. Breese, L. Sazanov, A universal coupling mechanism of respiratory complex I. Nature 609, 808–814 (2022).

9. D. Kampjut, L. A. Sazanov, The coupling mechanism of mammalian respiratory complex I. Science 370 (2020).

10. A. A. Stuchebrukhov, T. Hayashi, Single protonation of the reduced quinone in respiratory complex I drives four-proton pumping. FEBS Lett. 597, 237–245 (2023).

11. K. Parey, J. Lasham, D. J. Mills, A. Djurabekova, O. Haapanen, E. G. Yoga, H. Xie, W. Kühlbrandt, V. Sharma, J. Vonck, V. Zickermann, High-resolution structure and dynamics of mitochondrial complex I— Insights into the proton pumping mechanism. Sci. Adv. 7, eabj3221 (2021).

12. W. Fisher, J. J. Wright, M. N. Choy, G. M. Arantes, R. A. Waddell, D. N. Grba, J. Hirst, Conformational gating at histidine junctions coordinates proton translocation in respiratory complex I. [Preprint] (2026). 10.64898/2026.08.12.744383.

13. J. Gu, T. Liu, R. Guo, L. Zhang, M. Yang, The coupling mechanism of mammalian mitochondrial complex I. Nat. Struct. Mol. Biol. 29, 172–182 (2022).

14. A. Beghiah, P. Saura, S. Badolato, H. Kim, J. Zipf, D. Auman, A. P. Gamiz-Hernandez, J. Berg, G. Kemp, V. R. I. Kaila, Dissected antiporter modules establish minimal proton-conduction elements of the respiratory complex I. Nat. Commun. 15 (2024).

15. S. Badolato, S. C. Rossmann, J. Pereira, H. Kim, V. R. I. Kaila, Molecular Principles of Gating Proton Transport in the Antiporter Modules of Respiratory Complex I. J. Am. Chem. Soc., jacs.6c05956 (2026).

16. P. Saura, S. Badolato, A. Gangdal, A. Beghiah, V. R. I. Kaila, Disease-mutations perturb proton transfer reactions in respiratory complex I. Biochim. Biophys. Acta BBA - Bioenerg. 1867, 149602 (2026).

17. S. Dröse, S. Krack, L. Sokolova, K. Zwicker, H.-D. Barth, N. Morgner, H. Heide, M. Steger, E. Nübel, V. Zickermann, S. Kerscher, B. Brutschy, M. Radermacher, U. Brandt, Functional Dissection of the Proton Pumping Modules of Mitochondrial Complex I. PLoS Biol. 9, e1001128 (2011).

18. A. Di Luca, A. P. Gamiz-Hernandez, V. R. I. Kaila, Symmetry-related proton transfer pathways in respiratory complex I. Proc. Natl. Acad. Sci. U. S. A. 114, E6314–E6321 (2017).

19. M. E. Mühlbauer, P. Saura, F. Nuber, A. Di Luca, T. Friedrich, V. R. I. Kaila, Water-Gated Proton Transfer Dynamics in Respiratory Complex I. J. Am. Chem. Soc. 142, 13718–13728 (2020).

20. M. Röpke, P. Saura, D. Riepl, M. C. Pöverlein, V. R. I. Kaila, Functional Water Wires Catalyze Long-Range Proton Pumping in the Mammalian Respiratory Complex I. J. Am. Chem. Soc. 142, 21758–21766 (12 30).

21. F. Hoeser, P. Saura, C. Harter, V. R. I. Kaila, T. Friedrich, A leigh syndrome mutation perturbs long-range energy coupling in respiratory complex I. Chem. Sci. 16, 7374–7386 (2025).

22. C. Pecorilla, A. Altmeyer, O. Haapanen, Y. Lee, V. Zickermann, V. Sharma, Conformational dynamics of a histidine molecular switch in a cation/proton antiporter. Biochim. Biophys. Acta BBA - Bioenerg. 1866, 149563 (2025).

23. E. Endres, M. Torabi, M. Jousmäki, K. V. Huynh, C. Pecorilla, O. Zdorevskyi, V. Zickermann, V. Sharma, Molecular and energetic basis of histidine switch dynamics in respiratory complex I. Protein Sci. 35, e70720 (2026).

24. J. Liebau, D. Lazzaretti, T. Fürtges, A. Bichler, M. Pilsl, T. Rudack, R. Sprangers, 4D structural biology– quantitative dynamics in the eukaryotic RNA exosome complex. Nat. Commun. 16, 7896 (2025).

25. A. M. Gronenborn, Small, but powerful and attractive: 19F in biomolecular NMR. Structure 30, 6–14 (2022).

26. D. Rose-Sperling, M. A. Tran, L. M. Lauth, B. Goretzki, U. A. Hellmich, ^19^ F NMR as a versatile tool to study membrane protein structure and dynamics. Biol. Chem. 400, 1277–1288 (2019).

27. U. A. Hellmich, N. Pfleger, C. Glaubitz, ^19^ F-MAS NMR on Proteorhodopsin: Enhanced Protocol for Site-Specific Labeling for General Application to Membrane Proteins^†^. Photochem. Photobiol. 85, 535–539 (2009).

28. D. N. Grba, J. Hirst, Mitochondrial complex I structure reveals ordered water molecules for catalysis and proton translocation. Nat. Struct. Mol. Biol. 27, 892–900 (2020).

29. A. Beghiah, P. Saura, T. Kovalova, F. Hoeser, T. Friedrich, V. R. I. Kaila, A carboxylate switch point controls long-range energy transduction in respiratory Complex I. Nat. Commun. 17, 5737 (2026).

30. O. Haapanen, V. Sharma, Role of water and protein dynamics in proton pumping by respiratory complex I. Sci. Rep. 7, 7747 (2017).

31. G. Lipari, A. Szabo, Model-free approach to the interpretation of nuclear magnetic resonance relaxation in macromolecules. 1. Theory and range of validity. J. Am. Chem. Soc. 104, 4546–4559 (1982).

32. J. C. B. Dietschreit, A. Wagner, T. A. Le, P. Klein, H. Schindelin, T. Opatz, B. Engels, U. A. Hellmich, C. Ochsenfeld, Predicting^19^ F NMR Chemical Shifts: A Combined Computational and Experimental Study of a Trypanosomal Oxidoreductase–Inhibitor Complex. Angew. Chem. Int. Ed. 59, 12669–12673 (2020).

33. D. Flaig, M. Maurer, M. Hanni, K. Braunger, L. Kick, M. Thubauville, C. Ochsenfeld, Benchmarking Hydrogen and Carbon NMR Chemical Shifts at HF, DFT, and MP2 Levels. J. Chem. Theory Comput. 10, 572– 578 (2014).

34. T. W. Keal, D. J. Tozer, The exchange-correlation potential in Kohn–Sham nuclear magnetic resonance shielding calculations. J. Chem. Phys. 119, 3015–3024 (2003).

35. F. Jensen, Segmented Contracted Basis Sets Optimized for Nuclear Magnetic Shielding. J. Chem. Theory Comput. 11, 132–138 (2015).

36. F. Jensen, Basis Set Convergence of Nuclear Magnetic Shielding Constants Calculated by Density Functional Methods. J. Chem. Theory Comput. 4, 719–727 (2008).

37. V. Kaila, M. Wikstrom, G. Hummer, Electrostatics, hydration, and proton transfer dynamics in the membrane domain of respiratory complex I. Proc. Natl. Acad. Sci. U. S. A. 111, 6988–6993 (2014).

38. Y. Lee, O. Haapanen, A. Altmeyer, W. Kühlbrandt, V. Sharma, V. Zickermann, Ion transfer mechanisms in Mrp-type antiporters from high resolution cryoEM and molecular dynamics simulations. Nat. Commun. 13, 6091 (2022).

39. R. A. Waddell, H. L. Messent, M. M. Costa-Lima, J. J. Wright, J. H. Salisbury, J. Hirst, A modular mechanism determines the proton-pumping stoichiometry of respiratory complex I. [Preprint] (2026). 10.64898/2026.10.08.757549.

40. V. R. I. Kaila, Molecular Mechanisms Underlying Long-Range Proton-Coupled Electron Transfer and Energy Transduction in Complex I. *Chem. Rev.*, (in review) (2026).

41. J. M. Schuller, P. Saura, J. Thiemann, S. K. Schuller, A. P. Gamiz-Hernandez, G. Kurisu, M. M. Nowaczyk, V. R. I. Kaila, Redox-coupled proton pumping drives carbon concentration in the photosynthetic complex I. Nat Commun 11, 494 (1AD).

42. Z. Zhang, M. Zhang, R. L. Burnap, Action at a distance: The remarkable coupling of CO_2_ uptake to electron transfer in specialized cyanobacterial NDH-1 complexes. Proc. Natl. Acad. Sci. 122, e2511786122 (2025).

43. G. L. Chadwick, J. Hemp, W. W. Fischer, V. J. Orphan, Convergent evolution of unusual complex I homologs with increased proton pumping capacity: energetic and ecological implications. ISME J. 12, 2668–2680 (2018).

44. V. R. I. Kaila, M. I. Verkhovsky, M. Wikstrom, Proton-Coupled Electron Transfer in Cytochrome Oxidase. Chem. Rev. 110, 7062–7081 (2010).

45. M. Wikström, V. Sharma, V. Kaila, J. Hosler, G. Hummer, New Perspectives on Proton Pumping in Cellular Respiration. Chem. Rev. 115, 2196–2221 (2015).

46. P. Saura, D. Riepl, D. M. Frey, M. Wikström, V. R. I. Kaila, Electric fields control water-gated proton transfer in cytochrome *c* oxidase. Proc. Natl. Acad. Sci. U. S. A. 119, e2207761119 (2022).

47. M. Wikström, M. Verkhovsky, G. Hummer, Water-gated mechanism of proton translocation by cytochrome c oxidase. Biochim. Biophys. Acta-Bioenerg. 1604, 61–65 (2003).

48. R. Liang, J. Swanson, M. Wikstrom, G. Voth, Understanding the essential proton-pumping kinetic gates and decoupling mutations in cytochrome c oxidase. Proc. Natl. Acad. Sci. U. S. A. 114, 5924–5929 (2017).

49. R. Liang, J. Swanson, Y. Peng, M. Wikstrom, G. Voth, Multiscale simulations reveal key features of the proton-pumping mechanism in cytochrome c oxidase. Proc. Natl. Acad. Sci. U. S. A. 113, 7420–7425 (2016).

50. S. Jo, T. Kim, V. G. Iyer, W. Im, CHARMM-GUI: A web-based graphical user interface for CHARMM. J. Comput. Chem. 29, 1859–1865 (2008).

51. R. Best, X. Zhu, J. Shim, P. Lopes, J. Mittal, M. Feig, A. MacKerell, Optimization of the Additive CHARMM All-Atom Protein Force Field Targeting Improved Sampling of the Backbone phi, psi and Side-Chain chi(1) and chi(2) Dihedral Angles. J. Chem. Theory Comput. 8, 3257–3273 (2012).

52. J. C. Phillips, D. J. Hardy, J. D. C. Maia, J. E. Stone, J. V. Ribeiro, R. C. Bernardi, R. Buch, G. Fiorin, J. Hénin, W. Jiang, R. McGreevy, M. C. R. Melo, B. K. Radak, R. D. Skeel, A. Singharoy, Y. Wang, B. Roux, A. Aksimentiev, Z. Luthey-Schulten, L. V. Kalé, K. Schulten, C. Chipot, E. Tajkhorshid, Scalable molecular dynamics on CPU and GPU architectures with NAMD. J. Chem. Phys. 153, 044130 (2020).

53. S. G. Balasubramani, G. P. Chen, S. Coriani, M. Diedenhofen, M. S. Frank, Y. J. Franzke, F. Furche, R. Grotjahn, M. E. Harding, C. Hättig, A. Hellweg, B. Helmich-Paris, C. Holzer, U. Huniar, M. Kaupp, A. Marefat Khah, S. Karbalaei Khani, T. Müller, F. Mack, B. D. Nguyen, S. M. Parker, E. Perlt, D. Rappoport, K. Reiter, S. Roy, M. Rückert, G. Schmitz, M. Sierka, E. Tapavicza, D. P. Tew, C. van Wüllen, V. K. Voora, F. Weigend, A. Wodyński, J. M. Yu, TURBOMOLE: Modular program suite for *ab initio* quantum-chemical and condensed-matter simulations. J. Chem. Phys. 152, 184107 (2020).

54. W. Humphrey, A. Dalke, K. Schulten, VMD: Visual molecular dynamics. J. Mol. Graph. 14, 33–38 (1996).

55. E. C. Meng, T. D. Goddard, E. F. Pettersen, G. S. Couch, Z. J. Pearson, J. H. Morris, T. E. Ferrin, UCSF C_HIMERA_X: Tools for structure building and analysis. Protein Sci. 32, e4792 (2023).

56. N. Michaud-Agrawal, E. J. Denning, T. B. Woolf, O. Beckstein, MDAnalysis: A toolkit for the analysis of molecular dynamics simulations. J. Comput. Chem. 32, 2319–2327 (2011).

57. A. D. Becke, Density-functional thermochemistry. III. The role of exact exchange. J. Chem. Phys. 98, 5648– 5652 (1993).

58. C. Lee, W. Yang, R. G. Parr, Development of the Colle-Salvetti correlation-energy formula into a functional of the electron density. Phys. Rev. B 37, 785–789 (1988).

59. S. Grimme, J. Antony, S. Ehrlich, H. Krieg, A consistent and accurate *ab initio* parametrization of density functional dispersion correction (DFT-D) for the 94 elements H-Pu. J. Chem. Phys. 132, 154104 (2010).

60. A. Schäfer, H. Horn, R. Ahlrichs, Fully optimized contracted Gaussian basis sets for atoms Li to Kr. J. Chem. Phys. 97, 2571–2577 (1992).

61. C. I. Bayly, P. Cieplak, W. Cornell, P. A. Kollman, A well-behaved electrostatic potential based method using charge restraints for deriving atomic charges: the RESP model. J. Phys. Chem. 97, 10269–10280 (1993).

62. Y. Chen, B. Roux, Constant-pH Hybrid Nonequilibrium Molecular Dynamics–Monte Carlo Simulation Method. J. Chem. Theory Comput. 11, 3919–3931 (2015).

63. B. K. Radak, C. Chipot, D. Suh, S. Jo, W. Jiang, J. C. Phillips, K. Schulten, B. Roux, Constant-pH Molecular Dynamics Simulations for Large Biomolecular Systems. J. Chem. Theory Comput. 13, 5933–5944 (2017).

64. H. A. Stern, Molecular simulation with variable protonation states at constant pH. J. Chem. Phys. 126, 164112 (2007).

65. N. A. Baker, D. Sept, S. Joseph, M. J. Holst, J. A. McCammon, Electrostatics of nanosystems: Application to microtubules and the ribosome. Proc. Natl. Acad. Sci. U. S. A. 98, 10037–10041 (2001).

66. B. Rabenstein, G. M. Ullmann, E.-W. Knapp, Calculation of protonation patterns in proteins with structural relaxation and molecular ensembles - application to the photosynthetic reaction center. Eur. Biophys. J. 27, 626–637 (1998).

67. T. Meyer, G. Kieseritzky, E. Knapp, Electrostatic p *K*_a_ computations in proteins: Role of internal cavities. Proteins Struct. Funct. Bioinforma. 79, 3320–3332 (2011).

68. G. Kieseritzky, E. W. Knapp, Improved p *K*_a_ prediction: Combining empirical and semimicroscopic methods. J. Comput. Chem. 29, 2575–2581 (2008).

69. B. Brooks, C. Brooks, A. Mackerell, L. Nilsson, R. Petrella, B. Roux, Y. Won, G. Archontis, C. Bartels, S. Boresch, A. Caflisch, L. Caves, Q. Cui, A. Dinner, M. Feig, S. Fischer, J. Gao, M. Hodoscek, W. Im, K. Kuczera, T. Lazaridis, J. Ma, V. Ovchinnikov, E. Paci, R. Pastor, C. Post, J. Pu, M. Schaefer, B. Tidor, R. Venable, H. Woodcock, X. Wu, W. Yang, D. York, M. Karplus, CHARMM: The Biomolecular Simulation Program. J. Comput. Chem. 30, 1545–1614 (2009).

70. S. Riahi, C. N. Rowley, The CHARMM-TURBOMOLE interface for efficient and accurate QM/MM molecular dynamics, free energies, and excited state properties. J. Comput. Chem. 35, 2076–2086 (2014).

71. D. E. Drew, G. Von Heijne, P. Nordlund, J.-W. L. De Gier, Green fluorescent protein as an indicator to monitor membrane protein overexpression in *Escherichia coli*. FEBS Lett. 507, 220–224 (2001).

72. D. Drew, S. Newstead, Y. Sonoda, H. Kim, G. Von Heijne, S. Iwata, GFP-based optimization scheme for the overexpression and purification of eukaryotic membrane proteins in Saccharomyces cerevisiae. Nat. Protoc. 3, 784–798 (2008).

73. A. Hjelm, S. Schlegel, T. Baumgarten, M. Klepsch, D. Wickström, D. Drew, J.-W. De Gier, “Optimizing E. coli-Based Membrane Protein Production Using Lemo21(DE3) and GFP-Fusions” in Membrane Biogenesis, D. Rapaport, J. M. Herrmann, Eds. (Humana Press, Totowa, NJ, 2013; https://link.springer.com/10.1007/978-1-62703-487-6_24)vol. 1033 of *Methods in Molecular Biology*, pp. 381–400.

74. J. Liebau, M. Tersa, B. Trastoy, J. Patrick, A. Rodrigo-Unzueta, F. Corzana, T. Sparrman, M. E. Guerin, L. Mäler, Unveiling the activation dynamics of a fold-switch bacterial glycosyltransferase by 19F NMR. J. Biol. Chem. 295, 9868–9878 (2020).

75. F. Delaglio, S. Grzesiek, Geerten W., Vuister, G., Zhu, J. Pfeifer, A. Bax, NMRPipe: A multidimensional spectral processing system based on UNIX pipes. J. Biomol. NMR 6 (1995).

76. H. M. McConnell, Reaction Rates by Nuclear Magnetic Resonance. J. Chem. Phys. 28, 430–431 (1958).

77. P. Vallurupalli, A. Sekhar, T. Yuwen, L. E. Kay, Probing conformational dynamics in biomolecules via chemical exchange saturation transfer: a primer. J. Biomol. NMR 67, 243–271 (2017).

78. A. Godoy-Hernandez, A. H. Asseri, A. J. Purugganan, C. Jiko, C. De Ram, H. Lill, M. Pabst, K. Mitsuoka, C. Gerle, D. Bald, D. G. G. McMillan, Rapid and Highly Stable Membrane Reconstitution by LAiR Enables the Study of Physiological Integral Membrane Protein Functions. ACS Cent. Sci. 9, 494–507 (2023).

