## Supporting Information for "Electrostatic control of conformational dynamics drives proton translocation in respiratory Complex I"

**This PDF file includes:**

Figs. S1 to S15
Tables S1 to S6

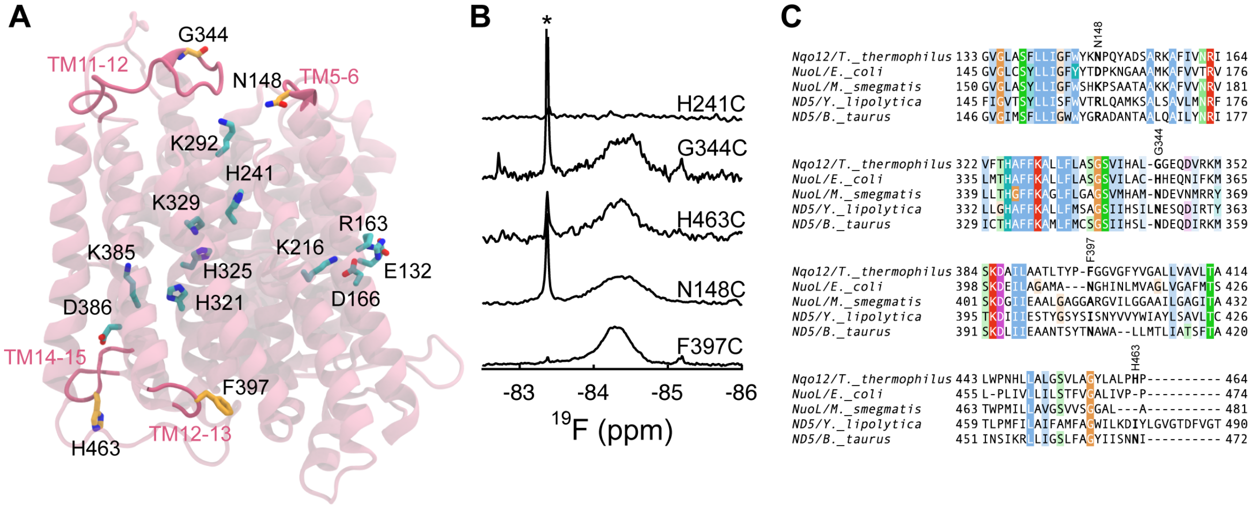


**Fig. S1. Design of $^{19}$F-labeled Nqo12 variants**. (**A**) Structure of Nqo12, showing functionally relevant residues (sidechains in cyan), and non-conserved residues for $^{19}$F-labeling (sidechains in yellow), located in loops exposed to the N-side (TM5-6 and TM11-12) or on the P-side (TM12-13 and TM14-15). (**B**) $^{19}$F-NMR spectra of the four labeled positions, indicated in panel A, acquired at 293 K. The asterisk indicates the resonance of the free TFA label. (**C**) Multiple sequence alignment (MSA), showing non-conserved residues at the selected label positions.

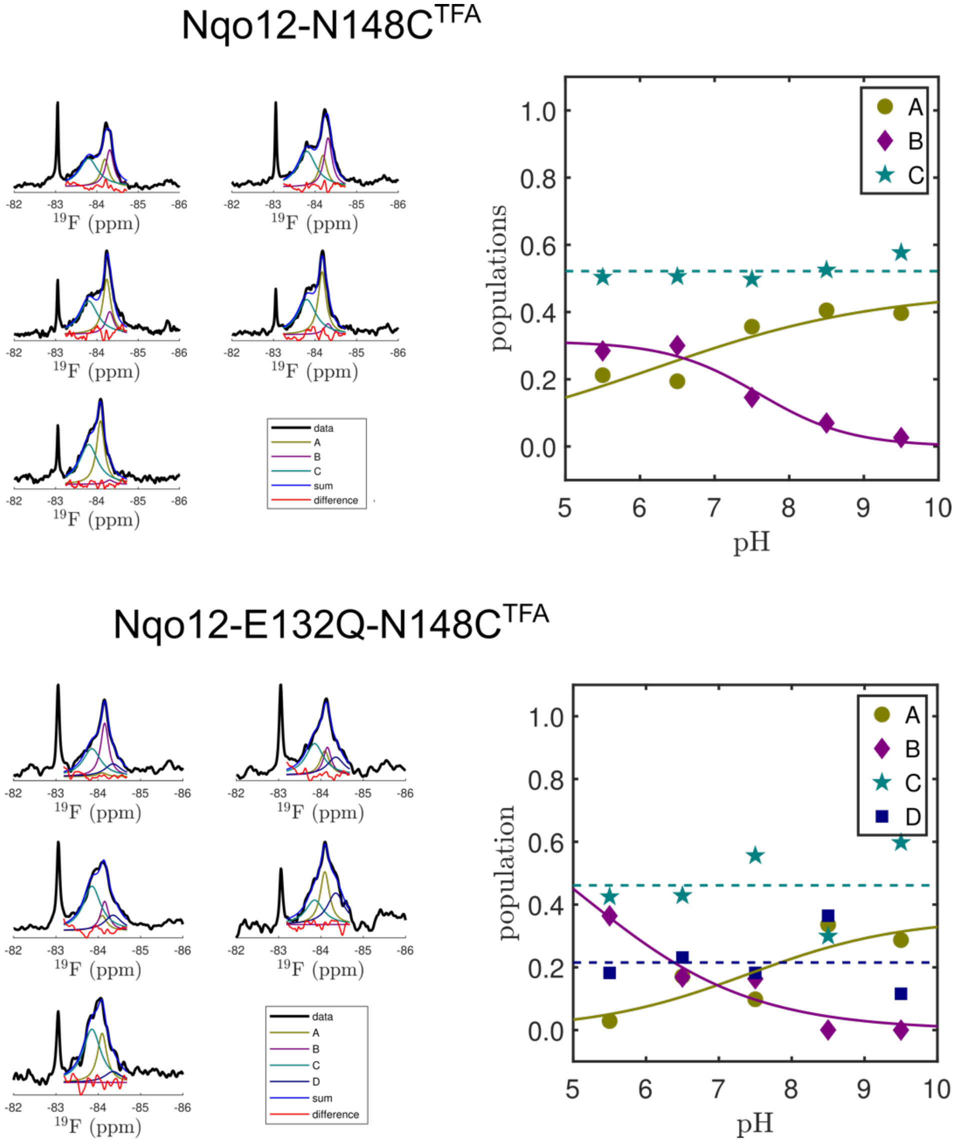


**Fig. S2. Line shape analysis and pH-dependence of observed NMR resonances.** Line shape analysis of $^{19}$F spectra at varying pH (*left*) and extracted populations (*right*) for WT-Nqo12 (*top*) and the E132Q-Nqo12 (*bottom*).

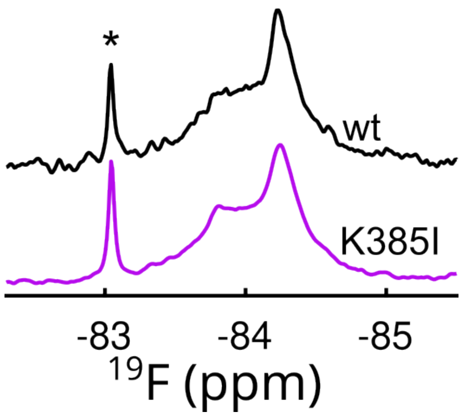


**Fig. S3. Effect of substitutions at the P-side proton pathway on $^{19}$F-NMR spectra.** The $^{19}$F-NMR spectrum is not affected by the K385I substitution at the P-side. The asterisk indicates the resonance of the free TFA label.

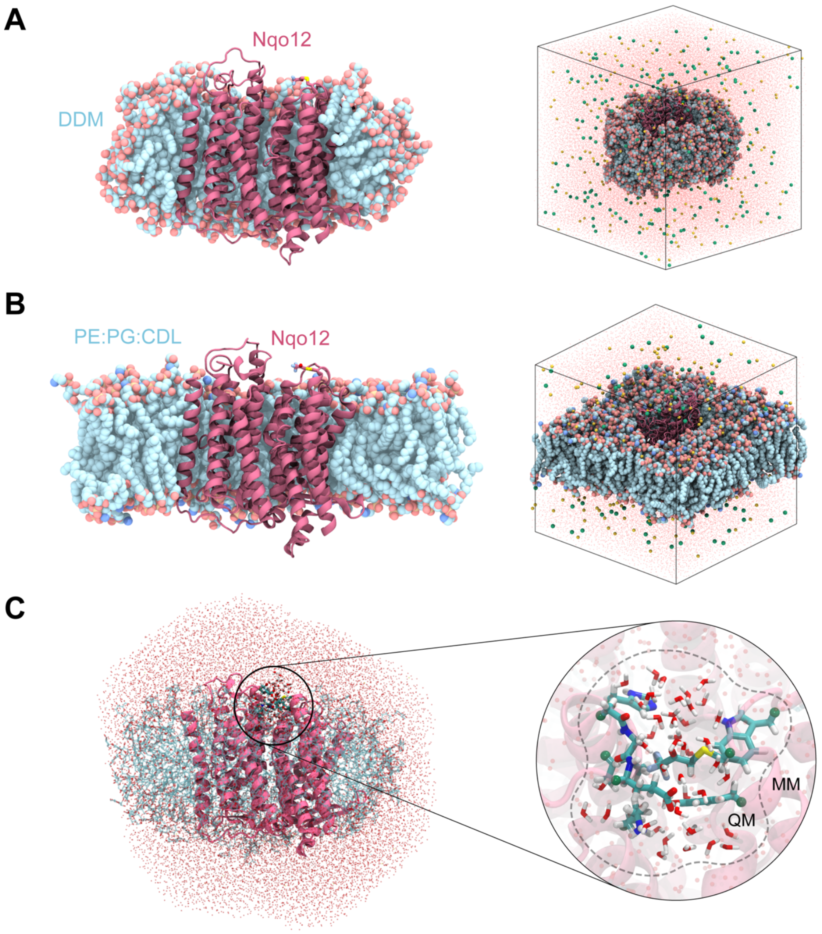


**Fig. S4. Setup for molecular dynamics and QM/MM simulations of the Nqo12 constructs.** Nqo12 embedded in (**A**) a DDM micelle, and (**B**) a POPE:POPG:CDL lipid membrane. (**C**) QM/MM setup of the TFA model in Nqo12.

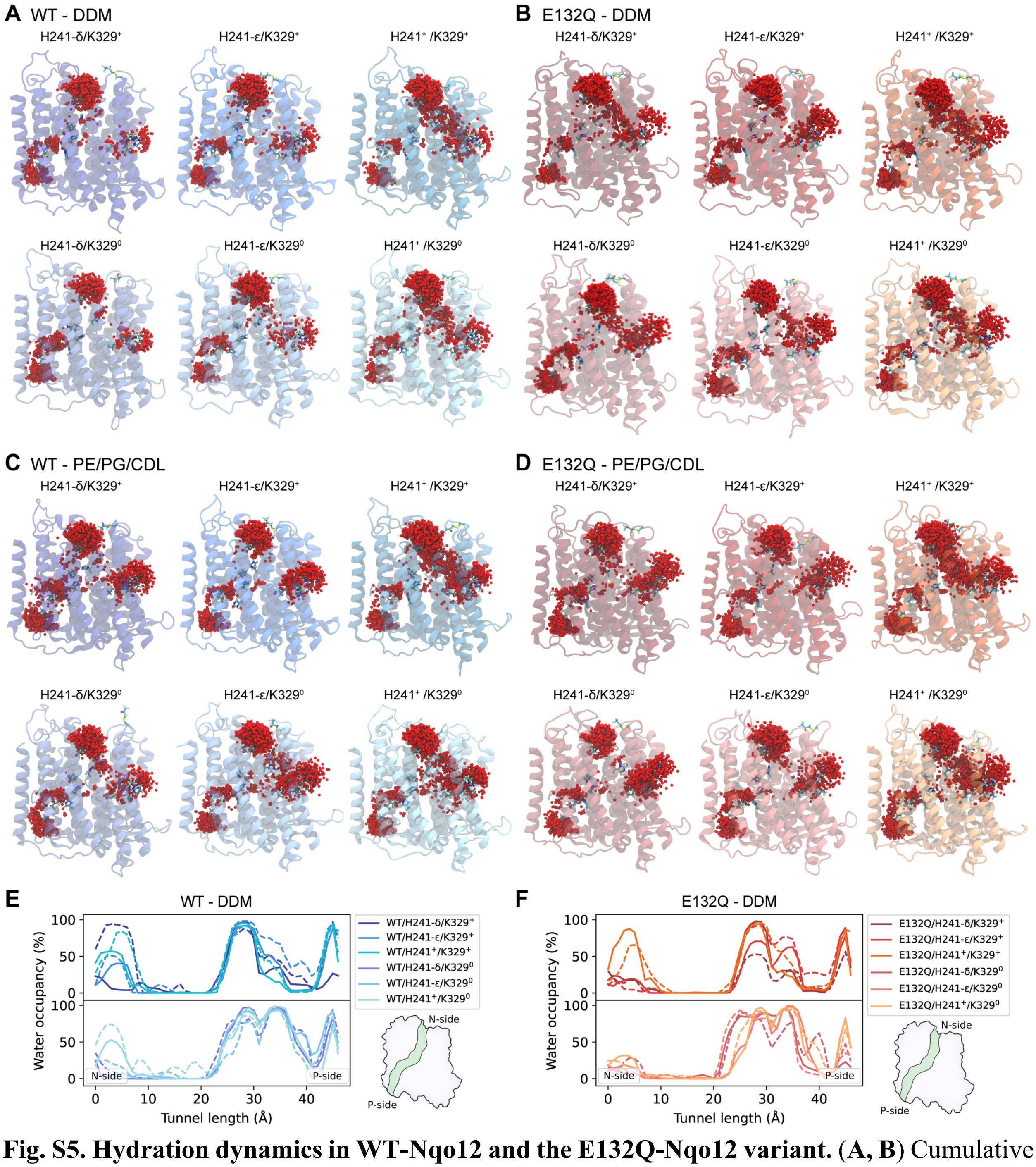


**Fig. S5. Hydration dynamics in WT-Nqo12 and the E132Q-Nqo12 variant.** (**A, B**) Cumulative hydration around key functional residues in (**A**) WT-Nqo12 and (**B**) the E132Q-Nqo12 variant for different protonation states of H241 and K329, from MD simulations carried out in DDM detergent. (**C, D**) Cumulative hydration around key functional residues in (**C**) WT-Nqo12 and (**D**) the E132Q-Nqo12 variant for different protonation states of H241 and K329, from MD simulations carried out in a PE:PG:CDL membrane. (**E, F**) Hydration along the N-side input pathway to P-side output pathway for (**E**) the WT-Nqo12 and (**F**) the E132Q-Nqo12 variant in different protonation states of H241 and K329, from MD simulations carried out in detergent.

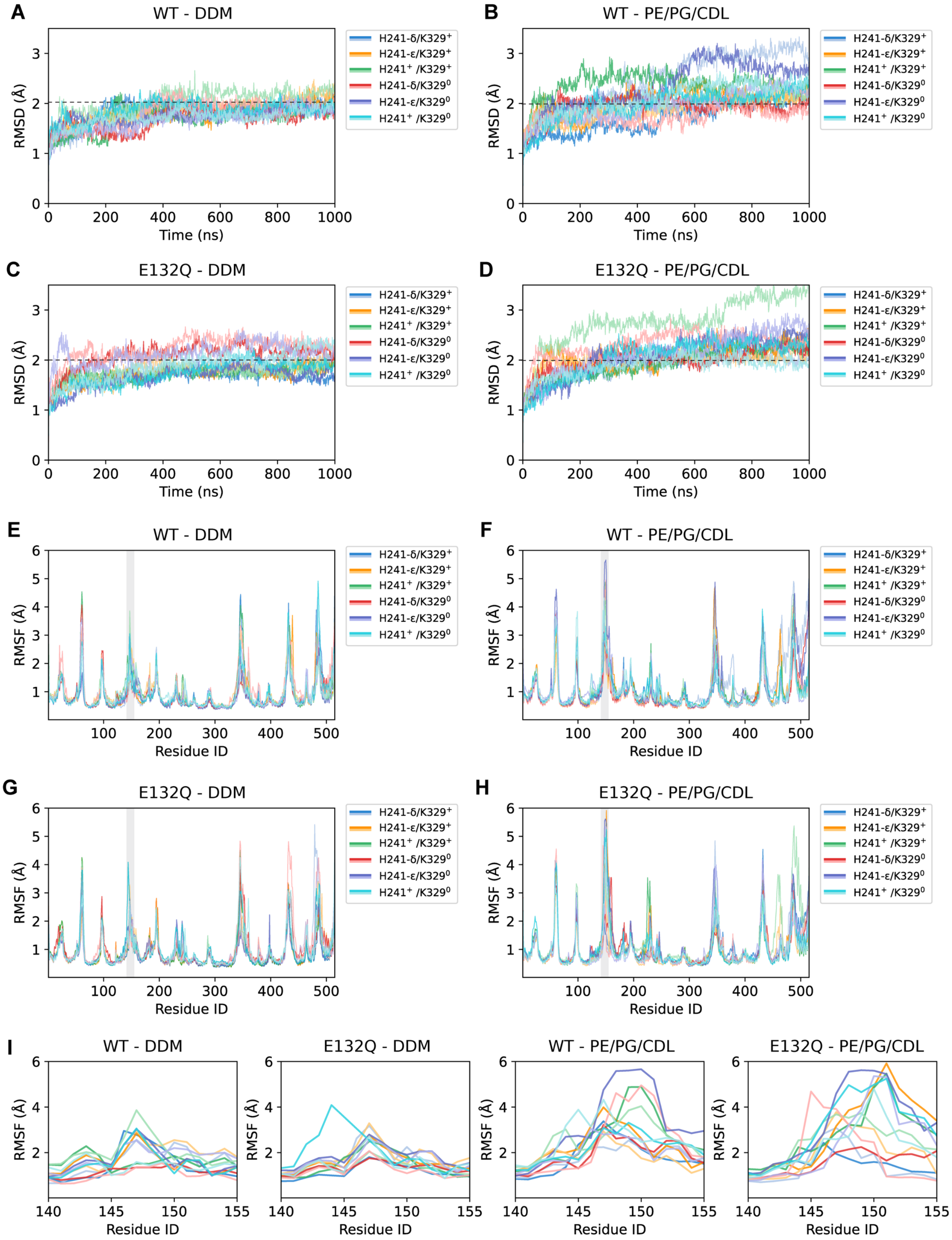


**Fig. S6. Dynamics of the WT-Nqo12 and the E132Q-Nqo12 variant in DDM detergent and membranes.** (**A-D**) *Root-mean-square-deviation* (RMSD) of the protein backbone in DDM and membranes. (**E-H**) *Root-mean-square-fluctuation* (RMSF) of protein Cα positions in DDM and membranes. (**I**) Close-up of the RMSF around the TM5-6 region, containing the $^{19}$F-label, suggesting that the region is more dynamic in the membrane simulations relative to detergents.

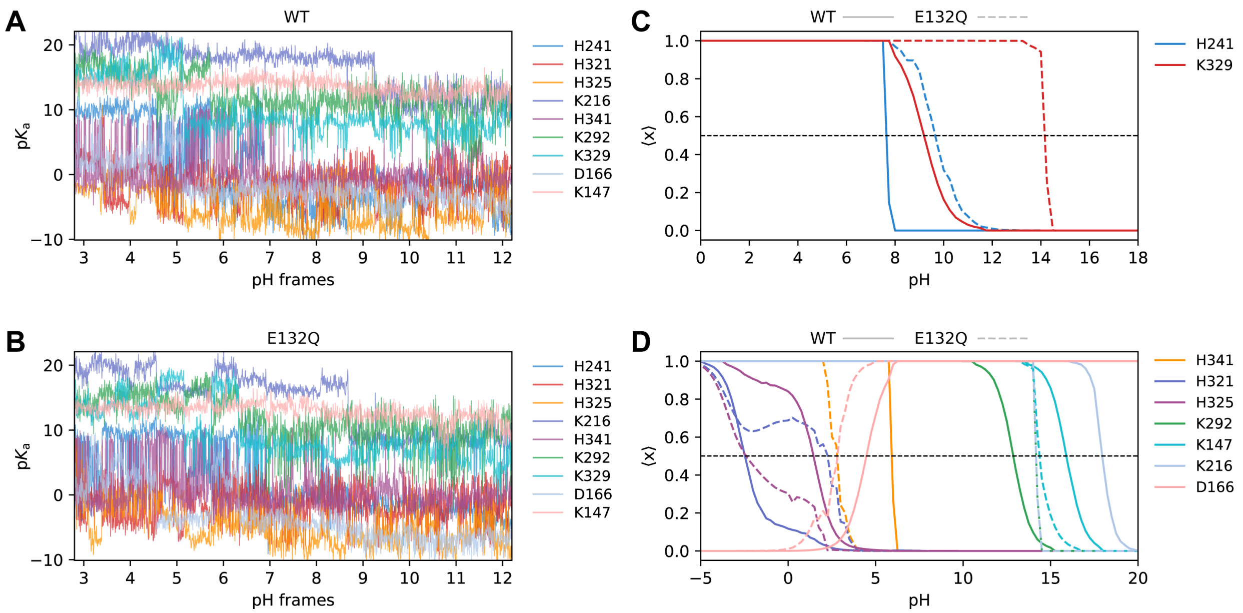


**Fig. S7. Constant pH-MD simulations of WT and E132Q-Nqo12. (A, B)** p$K_a$ of key residues along the cpH-MD trajectory at different pH values of (**A**) the WT and (**B**) the E132Q variant. (**C, D**) Titration curves of key residues along the proton pathway, showing the computed p$K_a$ of (**C**) H241 and K329 and (**D**) H341, H321, H325, K292, K147, K216, and D166 based on cpH-MD simulations and PBE/MC re-weighting (see *Methods*). The WT-Nqo12 model is shown in solid lines, and E132Q-Nqo12 in dashed lines.

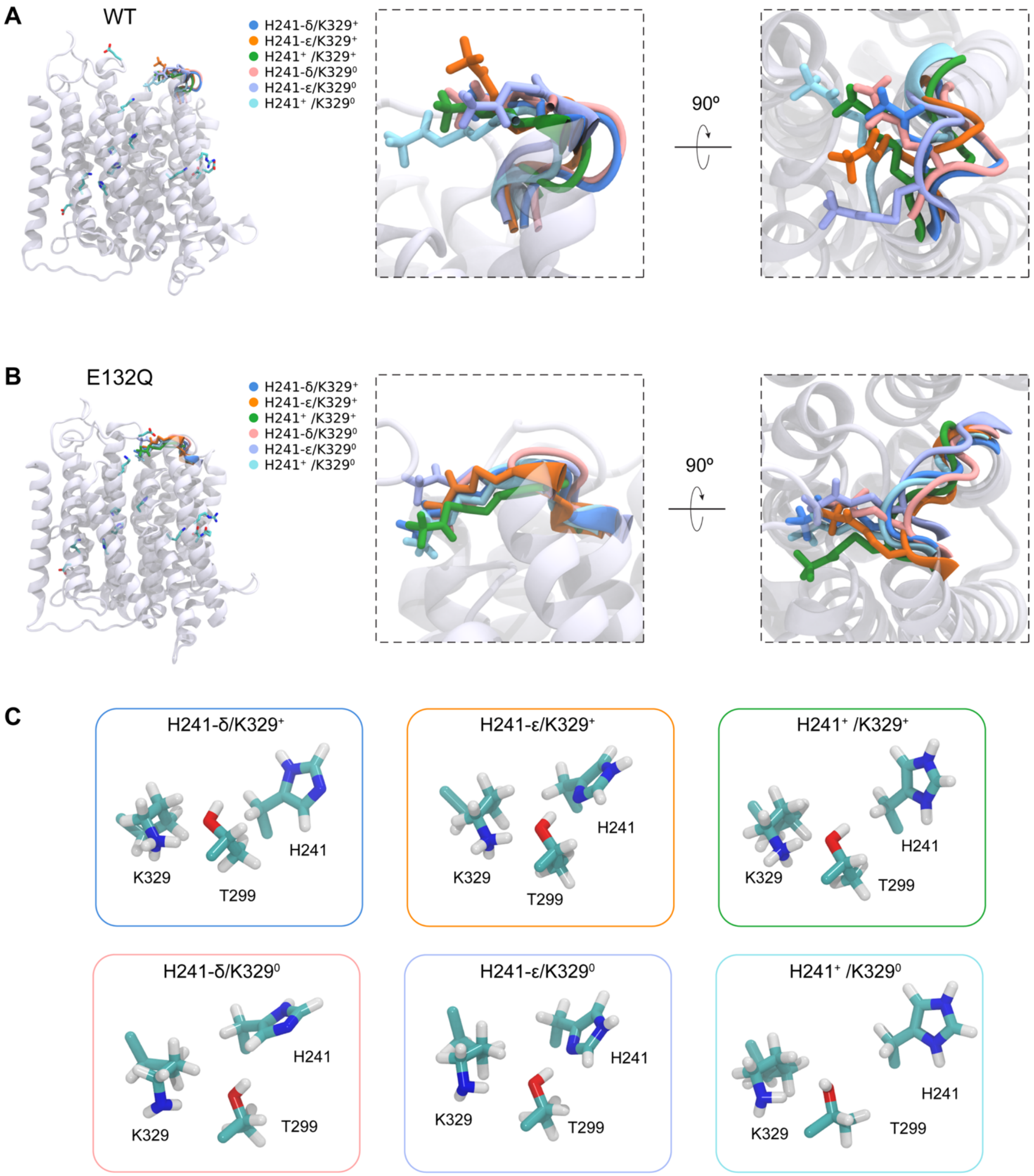


**Fig. S8. Conformational dynamics of the $^{19}$F-label based on clustering analysis.** (**A, B**) The figure shows the most populated cluster (cluster 1) in (**A**) WT-Nqo12 and (**B**) the E132Q-Nqo12 variant, for the different protonation states. (**C**) Conformational changes at the H241-K329-T299 interaction network in different protonation states.

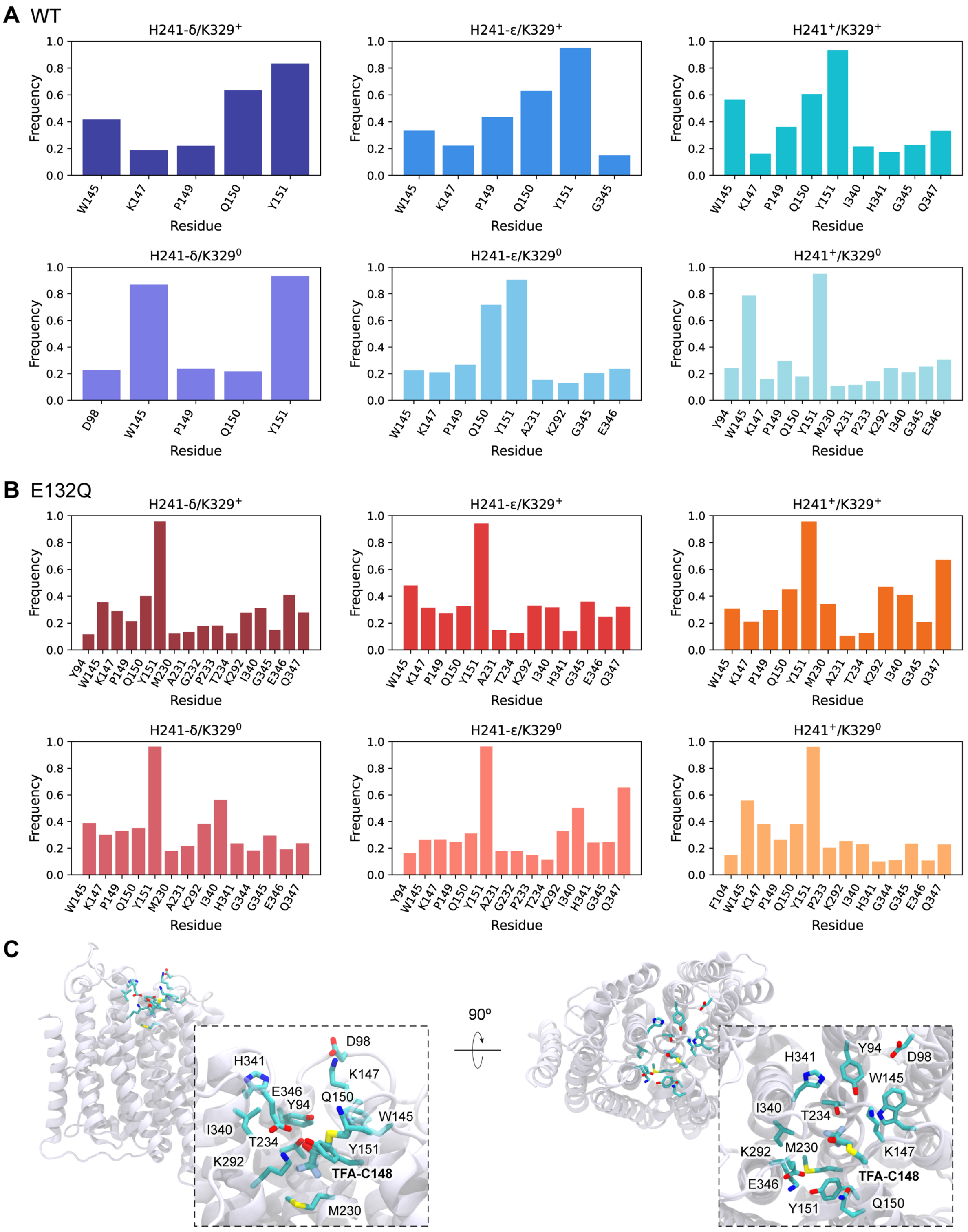


**Fig. S9. Interactions of the $^{19}$F-label based on contact analysis.** (**A**, **B**) Contact frequency of protein residues within < 5 Å of the $^{19}$F-label (excluding Cys148), with > 10% contact frequency during the MD simulations. The contact frequencies were calculated for two replicas for each protonation state (see Table S3). Residue contacts for (**A**) WT-Nqo12 and (**B**) E132Q-Nqo12. (**C**) Structural snapshot, showing key residues interacting with the $^{19}$F-label.

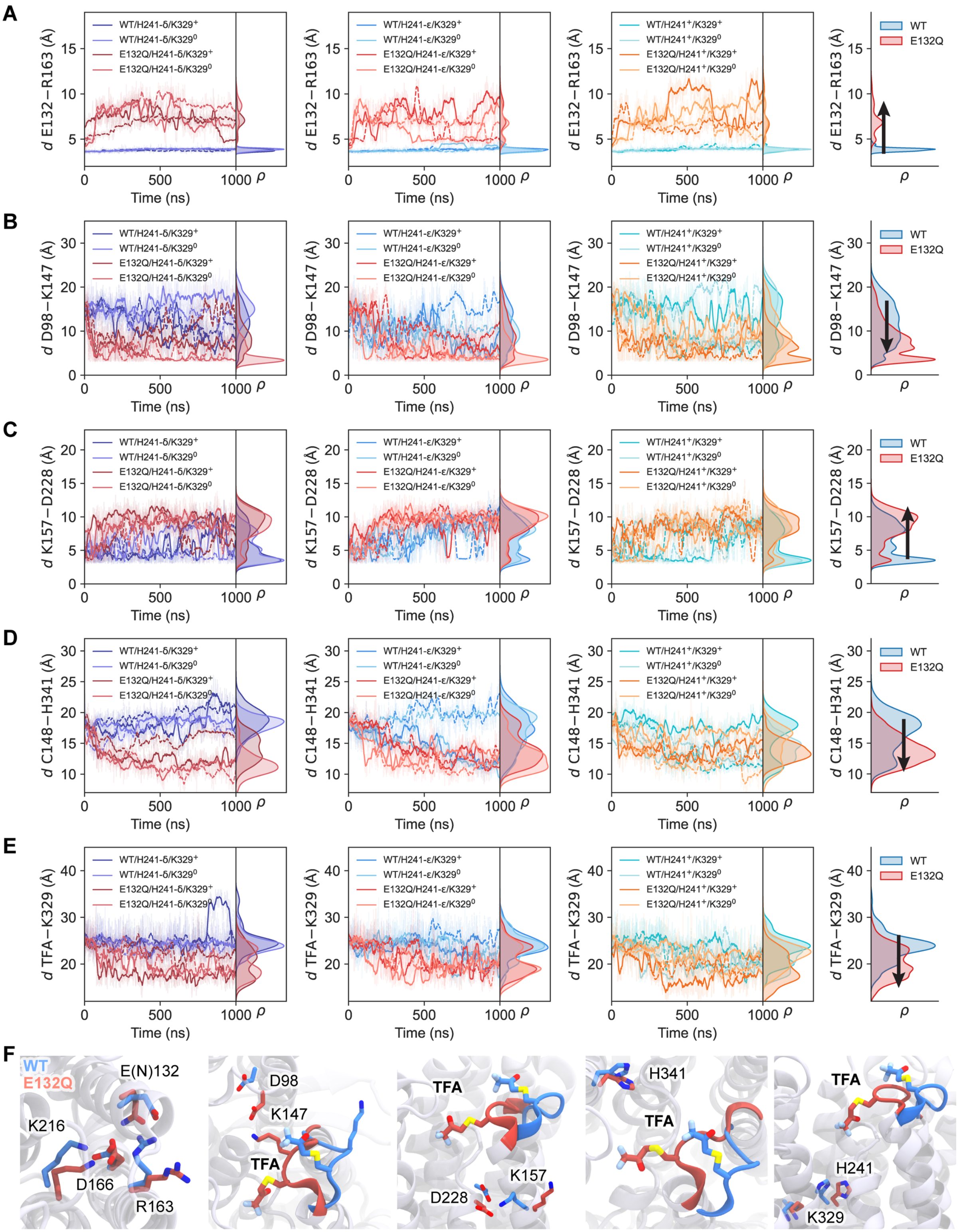


**Fig. S10. Distance analysis based on MD simulations of WT-Nqo12 and E132Q-Nqo12.** Distances during MD simulations of WT-Nqo12 and E132Q-Nqo12 for (**A**) the central ion-pair, E132-R163; (**B**) D98-K147 in the vicinity of the $^{19}$F-label; (**C**) K157-D228; (**D**) the $^{19}$F-label site C148-H341; (**E**) the $^{19}$F-label and K329. (**F**) Structural snapshots of MD simulations of the WT (in blue), and E132Q (in red).

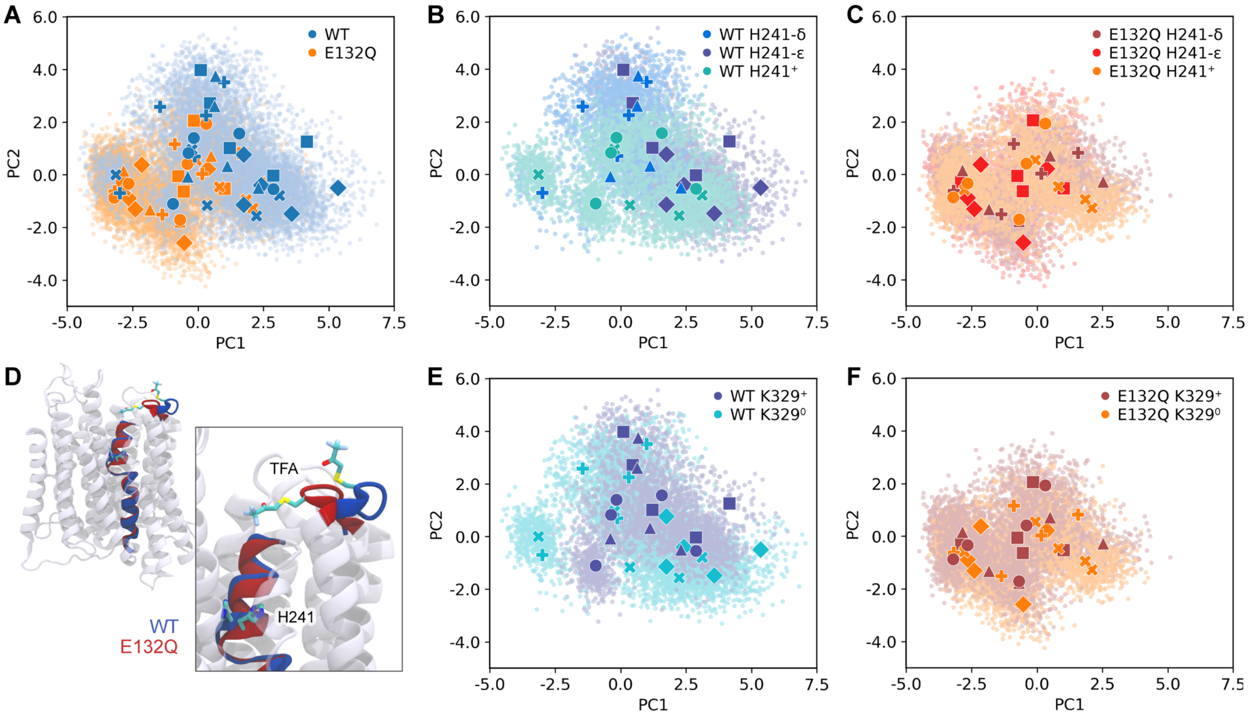


**Fig. S11. Principal component analysis of protonation-dependent global dynamics in the antiporter module Nqo12.** Principal components (PC) (**A**) PC1 and PC2 for all states, colored by the variant. The plot shows the WT (in blue) and E132Q (in orange), clustered into two distinct conformational ensembles. **(B, E)** PC1 and PC2 for all WT states, colored by the protonation state of (**B**) H241 and (**E**) K329. The dynamics of the protonated H241 state ($H241^{+}$, in green) overlaps with the conformations sampled by the E132Q-Nqo12 variant (see panel **A**). (**C**, **F**) PC1 and PC2 for all states of the E132Q-Nqo12 variant, colored by the protonation state of (**C**) H241 and (**F**) K329. **(D)** Structural snapshots of the H241 region, and position of the $^{19}$F-label in *inward* and *outward* conformations.

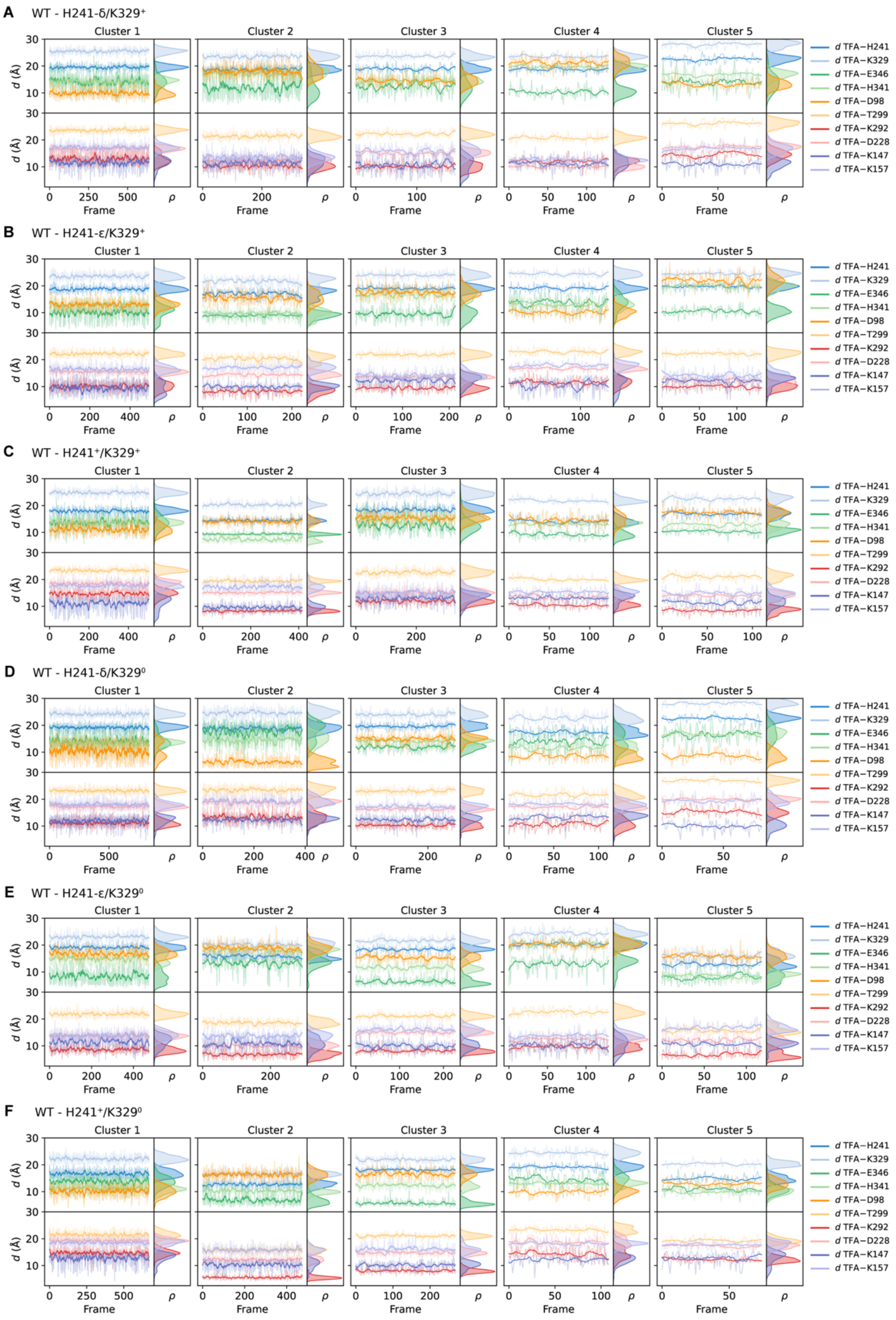


**Fig. S12. Selected distances from the $^{19}$F-label to key protein residues during MD simulations of WT-Nqo12.** (**A-F**) Central distances between the $^{19}$F-label and key protein residues in clusters 1 to 5 for WT-Nqo12.

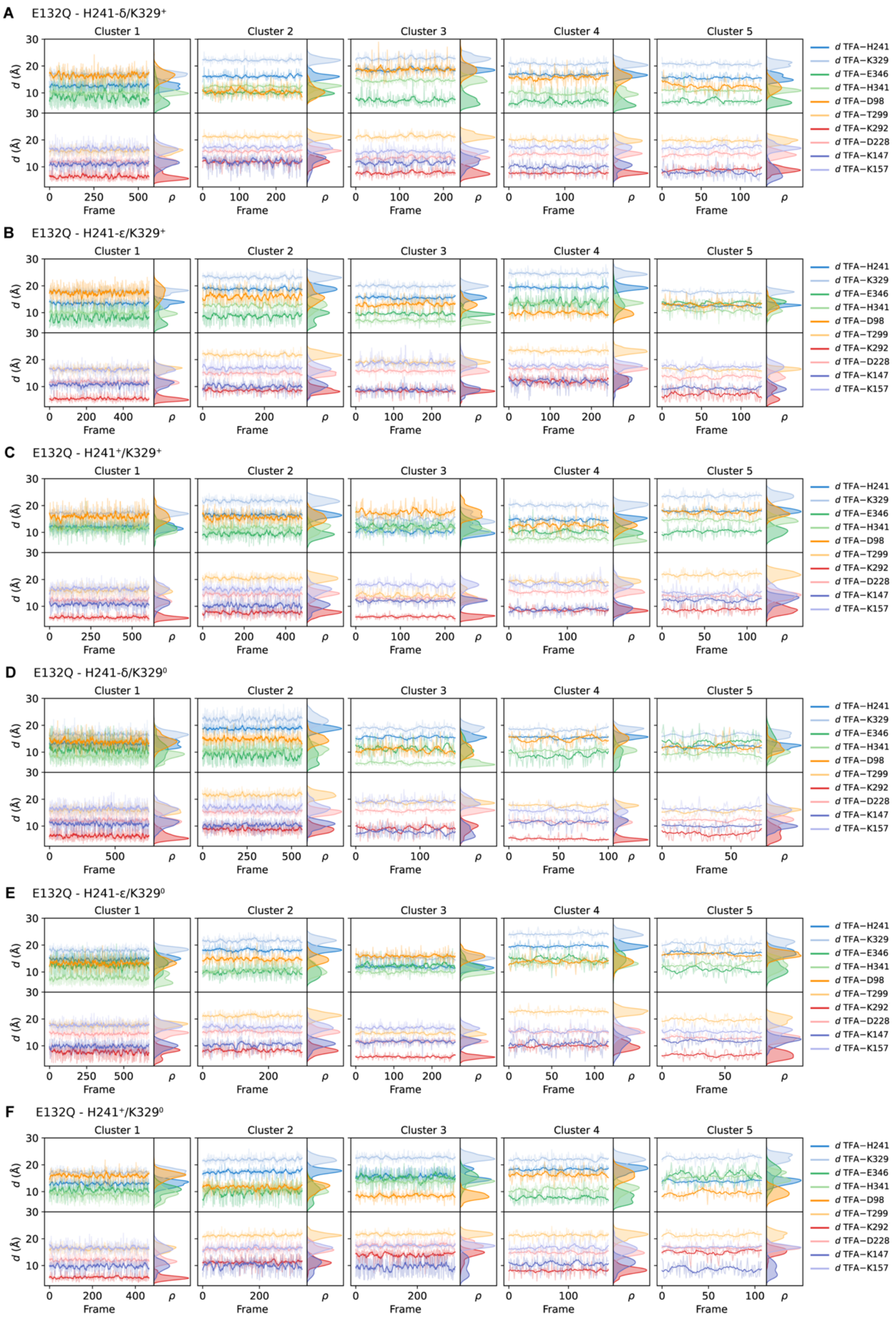


**Fig. S13. Selected distances from the [19]F-label to key protein residues during MD simulations of the E132Q-Nqo12 variant.** (**A-F**) Central distances between the [19]F-label and key protein residues in clusters 1 to 5 for the E132Q-Nqo12 variant.

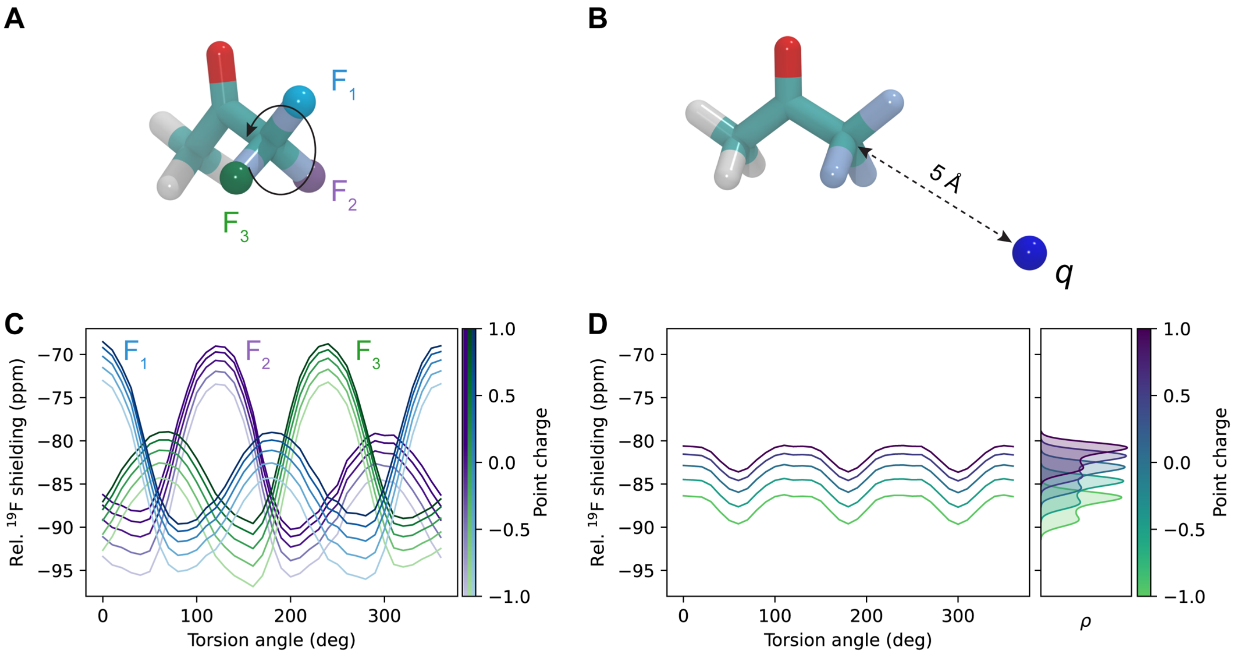


**Fig. S14. Electrostatic tuning of $^{19}$F chemical shifts of the TFA label based on DFT calculations.** (**A**) TFA model, showing the $CF_3$ rotation, and (**B**) the position of a point charge $q$, placed 5 Å from the $CF_3$ group of the TFA label, in the direction of the C-C bond. (**C**) $^{19}$F chemical shifts of the three fluorine atoms upon $CF_3$ rotation, calculated at the KT2/pcsSeg-2($CF_3$)/pcSseg-1(C,H,O)/MM level, combined with the tuning effect of the point charge $q$, with a charge ranging from -1$e$ (light colors) to +1$e$ (dark colors) (**D**) Shielding, averaged over the rotational angle, and its dependence on the charge ($q$), showing an electrostatic tuning of the chemical shifts.

.

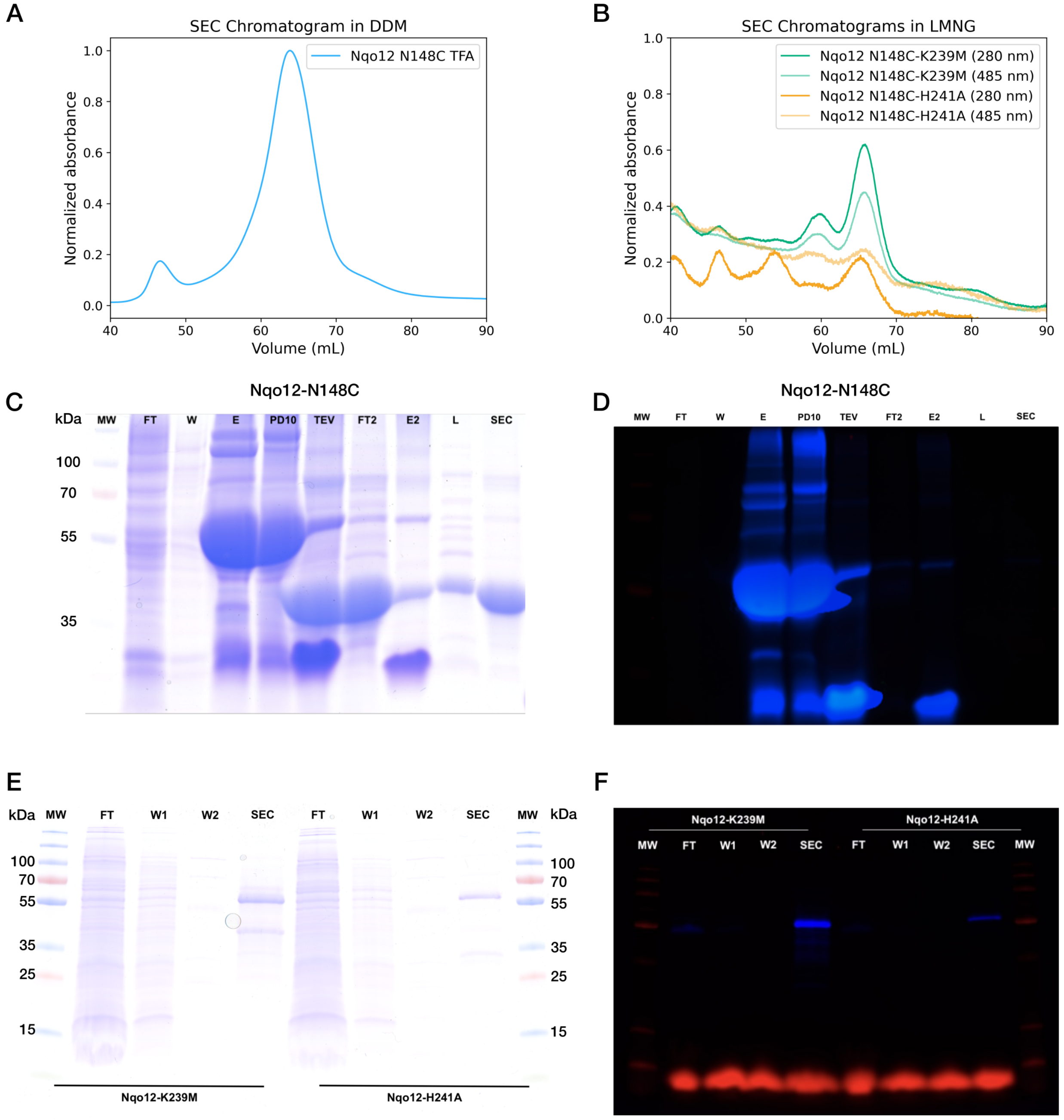


**Fig. S15. Protein expression and purification.** (**A**) SEC chromatogram of the Nqo12-N148C construct purified in DDM, showing normalized absorbance at 280 nm. (**B**) SEC chromatograms of the Nqo12-K329M and Nqo12-H241A variants, purified in LMNG, showing absorbances at 280 nm (protein) and 485 nm (sfGFP). (**C**) Coomassie gel and (**D**) in-gel fluorescence for Nqo12-N148C. (**E**) Coomassie gel and (**F**) in-gel fluorescence for Nqo12-K329M and Nqo12-H241A constructs. FT: flow through, W/W1/W2: wash steps, E: elution, PD10: sample after desalting, TEV: sample after TEV cleavage, FT2: flow through of inverse IMAC, E2: elution of inverse IMAC, L: sample after $^{19}$F-labeling, SEC: sample after size-exclusion chromatography.

**Table S1. Primer design for the antiporter constructs.**

| Construct | fwd primer | rev primer |
|---|---|---|
| Nqo12-N148C | CTTCTGGTACAAGTGTGCCCAGT ACGCCGACAG | CTTGTACCAGAAGCCGATGAGGAG |
| Nqo12-H241C | GTCTCCGCCCTGATCTGCGCGGC CACCATGGTGACGG | GATCAGGGCGGAGACGGGG |
| Nqo12-G344C | GTGATCCACGCCCTGTGTGGCGA GCAGGACGTGCGCAAG | CAGGGCGTGGATCACGCTCCC |
| Nqo12-H463C | TACCTGGCCCTGCCCTGTCCCTT GCCCAACGTCCTGGAACC | GGGCAGGGCCAGGTACCCCG |
| Nqo12-F397C | ACCCTCACCTACCCCTGTGGGGG AGTGGGGTTCTACGTGG | GGGGTAGGTGAGGGTGGCGGC |
| Nqo12-N148C-K385I | TCCGGCTTCTGGTCCATCGACGC CATCCTCGCCG | CCAGAAGCCGGAAAGCAAGGGG |
| Nqo12-N148C-E132Q | ATGTTCATCGGCTGGCAGGGGGT GGGCCTG | CCAGCCGATGAACATCACC |
| Nqo12-N148C-K329M | CCACGCCTTCTTCATGGCCCTCC TCTTCCTG | GAAGAAGGCGTGGGTGAAGACG |
| Nqo12-N148C-H241A | CTCCGCCCTGATCGCCGCGGCCA CCATG | GATCAGGGCGGAGACGGGGG |

**Table S2. Data fitting parameters for the CEST experiment of the Nqo12$^{N148C\text{-}TFA}$ construct.**

| Parameter | value |
|---|---|
| $k_{ex}$ | 1.9±0.8 $s^{-1}$ |
| $p_C$ | 0.29±0.19 |
| Δν | 0.49±0.03 ppm |
| $R_1$ | 4.1±0.3 $s^{-1}$ |
| $R_{2,A/B}$ | 347±41 $s^{-1}$ |
| $R_{2,C}$ | 750±166 $s^{-1}$ |

**Table S3. List of classical MD simulations.** Other residues are modeled in their standard protonation states, unless otherwise indicated (see *Methods*).

| **Simulation** | **System** | **State** | **Environment** | **Length (ns)** | **Replicas** |
|---|---|---|---|---|---|
| S1-S2 | WT | H241-δ/K329$^+$/K385$^0$ | DDM | 1000 | 2 |
| S3-S4 | WT | H241-ε/K329$^+$/K385$^0$ | DDM | 1000 | 2 |
| S5-S6 | WT | H241$^+$/K329$^+$/K385$^0$ | DDM | 1000 | 2 |
| S7-S8 | WT | H241-δ/K329$^0$/K385$^+$ | DDM | 1000 | 2 |
| S9-S10 | WT | H241-ε/K329$^0$/K385$^+$ | DDM | 1000 | 2 |
| S11-S12 | WT | H241$^+$/K329$^0$/K385$^+$ | DDM | 1000 | 2 |
| S13-S14 | E132Q | H241-δ/K329$^+$/K385$^0$ | DDM | 1000 | 2 |
| S15-S16 | E132Q | H241-ε/K329$^+$/K385$^0$ | DDM | 1000 | 2 |
| S17-S18 | E132Q | H241$^+$/K329$^+$/K385$^0$ | DDM | 1000 | 2 |
| S19-S20 | E132Q | H241-δ/K329$^0$/K385$^+$ | DDM | 1000 | 2 |
| S21-S22 | E132Q | H241-ε/K329$^0$/K385$^+$ | DDM | 1000 | 2 |
| S23-S24 | E132Q | H241$^+$/K329$^0$/K385$^+$ | DDM | 1000 | 2 |
| S25-S26 | WT | H241-δ/K329$^+$/K385$^0$ | PE/PG/CDL | 1000 | 2 |
| S27-S28 | WT | H241-ε/K329$^+$/K385$^0$ | PE/PG/CDL | 1000 | 2 |
| S29-S30 | WT | H241$^+$/K329$^+$/K385$^0$ | PE/PG/CDL | 1000 | 2 |
| S31-S32 | WT | H241-δ/K329$^0$/K385$^+$ | PE/PG/CDL | 1000 | 2 |
| S33-S34 | WT | H241-ε/K329$^0$/K385$^+$ | PE/PG/CDL | 1000 | 2 |
| S35-S36 | WT | H241$^+$/K329$^0$/K385$^+$ | PE/PG/CDL | 1000 | 2 |
| S37-S38 | E132Q | H241-δ/K329$^+$/K385$^0$ | PE/PG/CDL | 1000 | 2 |
| S39-S40 | E132Q | H241-ε/K329$^+$/K385$^0$ | PE/PG/CDL | 1000 | 2 |
| S41-S42 | E132Q | H241$^+$/K329$^+$/K385$^0$ | PE/PG/CDL | 1000 | 2 |
| S43-S44 | E132Q | H241-δ/K329$^0$/K385$^+$ | PE/PG/CDL | 1000 | 2 |
| S45-S46 | E132Q | H241-ε/K329$^0$/K385$^+$ | PE/PG/CDL | 1000 | 2 |
| S47-S48 | E132Q | H241$^+$/K329$^0$/K385$^+$ | PE/PG/CDL | 1000 | 2 |
| | | | **TOTAL:** | **48 µs** | |

**Table S4. Clustering analysis of the $^{19}$F-probe in MD simulations.** The five most populated clusters are reported, showing the percentage of frames belonging to each cluster, based on an RMSD cutoff of 2.5 Å. For each state, a total of 2000 frames were analyzed from two MD simulation replicas, including 2 μs of data (1 frame/ns).

| System | Cluster 1 | Cluster 2 | Cluster 3 | Cluster 4 | Cluster 5 | Unassigned |
|---|---|---|---|---|---|---|
| WT - H241-δ/K329$^+$ | 31.9 | 16.8 | 8.2 | 6.4 | 4.5 | 32.1 |
| WT - H241-δ/K329$^0$ | 41.9 | 19.4 | 13.8 | 5.6 | 4.1 | 15.1 |
| WT - H241-ε/K329$^+$ | 24.8 | 11.2 | 10.7 | 7.0 | 6.7 | 39.6 |
| WT - H241-ε/K329$^0$ | 23.8 | 14.7 | 11.4 | 6.5 | 6.0 | 37.7 |
| WT - H241$^+$/K329$^+$ | 25.2 | 20.8 | 16.5 | 6.2 | 5.6 | 25.8 |
| WT - H241$^+$/K329$^0$ | 33.6 | 23.8 | 13.4 | 5.4 | 3.8 | 20.0 |
| E132Q - H241-δ/K329$^+$ | 28.0 | 13.6 | 11.4 | 9.0 | 6.4 | 31.6 |
| E132Q - H241-δ/K329$^0$ | 38.0 | 28.0 | 7.8 | 5.0 | 3.6 | 17.6 |
| E132Q - H241-ε/K329$^+$ | 27.0 | 16.2 | 13.2 | 12.1 | 6.4 | 25.2 |
| E132Q - H241-ε/K329$^0$ | 33.6 | 15.2 | 13.1 | 5.8 | 4.0 | 28.4 |
| E132Q - H241$^+$/K329$^+$ | 30.2 | 24.0 | 11.2 | 8.6 | 5.9 | 20.2 |
| E132Q - H241$^+$/K329$^0$ | 23.2 | 17.6 | 16.2 | 8.4 | 5.4 | 29.2 |

**Table S5. List of QM/MM simulations.**

| System | State | Probe position | QM region (size) | Sampling |
|---|---|---|---|---|
| WT | H241-δ/ $K329^{+}/K385^{0}$ | Exposed | TFA, Cys148, Gln150, 77 water molecules ($N$ = 257 atoms) | 5 ps |
| WT | H241ε/ $K329^{+}/K385^{0}$ | Intermediate | TFA, Cys148, Trp145, Gln150, Tyr151, 56 water molecules ($N$=229 atoms) | 5 ps |
| WT | $H241^{+}$/ $K329^{0}/K385^{+}$ | Buried | TFA, Cys148, Tyr151, Trp145, Lys292, Ile340, His341, Glu346, Gly344-Gly345-Glu346 backbone, 47 water molecules ($N$=260 atoms) | 5 ps |
| E132Q | H241-ε/ $K329^{+}/K385^{0}$ | Buried | TFA, Cys148, Tyr151, Trp145, Lys292, Ile340, His341, Glu346, Gly344-Gly345-Glu346 backbone, 52 water molecules ($N$=275 atoms) | 5 ps |
| | | | **Total:** | 20 ps |

**Table S6. List of employed buffers.**

| Buffer | Function | Composition |
|---|---|---|
| A | Resuspension buffer | 25 mM HEPES pH 7.5, 150 mM NaCl |
| B | Wash buffer | 25 mM HEPES pH 7.5, 300 mM NaCl, 0.05% DDM, supplemented with 20, 75, 125, and 150 mM imidazole |
| C | Elution buffer | 25 mM HEPES pH 7.5, 300 mM NaCl, 0.05% DDM, 300 mM imidazole |
| D | SEC buffer, sample buffer | 25 mM HEPES pH 7.5, 300 mM NaCl, 0.05% DDM |
| pH 5.5 | Sample buffer | 25 mM MES pH 5.5, 300 mM NaCl, 0.05% DDM |
| pH 6.5 | Sample buffer | 25 mM MES pH 6.5, 300 mM NaCl, 0.05% DDM |
| pH 8.5 | Sample buffer | 25 mM Tris/HCl pH 8.5, 300 mM NaCl, 0.05% DDM |
| pH 9.5 | Sample buffer | 25 mM Tris/HCl pH 9.5, 300 mM NaCl, 0.05% DDM |
| E | Proteoliposome buffer | 2 mM HEPES/KOH, 50 mM KCl, 5 mM $MgCl_2$, pH 7.3 |
